# Phenotypic Pliancy and the Breakdown of Epigenetic Polycomb Mechanisms

**DOI:** 10.1101/2022.01.18.476783

**Authors:** Maryl Lambros, Yehonatan Sella, Aviv Bergman

**Affiliations:** Department of Systems and Computational Biology, Albert Einstein College of Medicine, Bronx, NY 10461, USA; Dominick P. Purpura Department of Neuroscience, Albert Einstein College of Medicine, Bronx, NY 10461, USA; Department of Pathology, Albert Einstein College of Medicine, Bronx, NY 10461, USA; Santa Fe Institute, Santa Fe, NM 87501, USA

## Abstract

Epigenetic regulatory mechanisms allow multicellular organisms to develop distinct specialized cell identities despite having the same total genome. Cell-fate choices are based on gene expression programs and environmental cues that cells experience during embryonic development, and are usually maintained throughout the life of the organism despite new environmental cues. The evolutionarily conserved Polycomb group (PcG) proteins form Polycomb Repressive Complexes that help orchestrate these developmental choices. Post-development, these complexes actively maintain the resulting cell fate, even in the face of environmental perturbations. Given the crucial role of these polycomb mechanisms in providing phenotypic fidelity (i.e. maintenance of cell fate), we hypothesize that their dysregulation after development will lead to decreased phenotypic fidelity allowing dysregulated cells to sustainably switch their phenotype in response to environmental changes. We call this abnormal phenotypic switching *phenotypic pliancy*. We introduce a general computational evolutionary model that allows us to test our systems-level phenotypic pliancy hypothesis *in-silico* and in a context-independent manner. We find that 1) phenotypic fidelity is an emergent systems-level property of PcG-like mechanism evolution, and 2) phenotypic pliancy is an emergent systems-level property resulting from this mechanism’s dysregulation. Since there is evidence that metastatic cells behave in a phenotypically pliant manner, we hypothesize that progression to metastasis is driven by the emergence of phenotypic pliancy in cancer cells as a result of PcG mechanism dysregulation. We corroborate our hypothesis using single-cell RNA-sequencing data from metastatic cancers. We find that metastatic cancer cells are phenotypically pliant in the same manner as predicted by our model.

**Significance Statement:** We introduce the concept of cellular phenotypic pliancy– sustained abnormal phenotypic switching in response to environmental changes– and demonstrate that such behavior can be caused by dysregulation of Polycomb mechanisms. To overcome the incomplete knowledge about these mechanisms in higher organisms, we develop an abstract computational model to study the emergence of phenotypic pliancy from a general systems-level view by confirming our hypothesis over a wide range of simulated gene-regulatory networks and Polycomb mechanism patterns. We corroborate our hypothesis and model predictions using single-cell RNA-seq metastatic cancer datasets. Our hypothesis has the potential to shed light on a general phenomenon for complex diseases where abnormal phenotypic switching is relevant.

## Introduction

During early development in multicellular organisms, the differentiated phenotype of cells, i.e. cell identity, is sensitive to and determined by the environmental influences [1, 2]. However, after development, cells tend to maintain their identity, becoming markedly less sensitive to environmental changes even as they may exhibit some limited phenotypic plasticity [3, 4]. We will refer to the maintenance of cellular identity post-development as *phenotypic fidelity*.

Because all cells in a multicellular organism possess the same genome, a key regulator of cell-fate choices and cell identity maintenance is epigenetic regulation mechanisms, which emerged during the expansion of the Metazoa [5–7]. One of the most prominent and enigmatic epigenetic regulatory mechanisms is the evolutionarily conserved polycomb mechanisms involving the Polycomb group (PcG) and Trithorax group (TrxG) proteins that form complexes (called Polycomb Repressive Complexes (PRCs) for PcG proteins) thought to have evolved at the dawn of multicellularity [8, 9]. PcG and TrxG proteins act antagonistically to cause heritable alterations in gene expression or function, without changes in DNA sequence. PcG proteins act by continually repressing a set of target genes if their expression, as influenced by the environment and maternal gene expression levels, falls below a certain threshold at a critical time point during development [3, 10–14]. Conversely, TrxG proteins act to counteract PcG-mediated repression and to continually maintain the memory of the transcriptional states of the target genes [14, 15]. The polycomb mechanism promotes cellular differentiation during development in response to environmental inputs, and *actively maintains* cell identity post-development even in the face of environmental perturbations (see Figure 1) [3, 9, 13–15]. It has also been demonstrated computationally that a PcG-like mechanism allows dynamic altering of a gene regulatory network to create multiple gene-regulatory sub-networks, resulting in different phenotypes from the same genotype(see Figure 1) [16].

**Fig 1.**
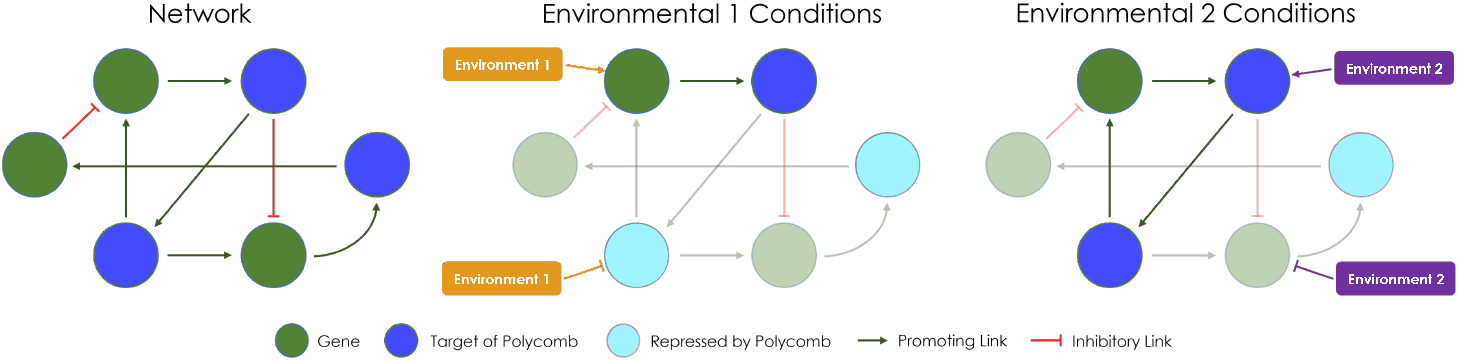
Schematic of Polycomb Mechanisms During Development: The evolution of Polycomb mechanisms carves the overall gene regulatory network into subnetworks based on the environments encountered during development, which leads to different differentiated phenotypes at the end of development.

Given the crucial role of polycomb mechanisms in providing phenotypic fidelity, we hypothesize that dysregulation of polycomb mechanisms after development will lead to decreased phenotypic fidelity, which will allow dysregulated cells to directly and sustainably switch their phenotype in response to environmental changes. We call this abnormal phenotypic switching in response to environmental changes *phenotypic pliancy*. Here, the environment encompasses both the external environment, such as availability of oxygen, temperature, and perhaps a local microbiome, and the signaling environment created by cellular communication networks. We further hypothesize that cells with appropriate dysregulated polycomb mechanisms, when introduced into a new environment will switch into a phenotype that more closely, *but not fully*, resembles that of the already evolved phenotype in this environment since we expect that the environmental cues will dominate once the cellular memory disappears. For example, a dysregulated breast cell placed in a lung environment would alter its phenotype to more resemble that of a lung cell than a breast cell if the PRC(s) dysregulated in the breast cell derepressed the target genes important for lung cell determination. Note, while phenotypic *plasticity* typically refers to naturally occurring variation in response to typical environmental changes, such that a breast cell’s phenotype will have some variation in response to the variation in the breast environment, phenotypic *pliancy* refers to changes in phenotype in response to drastic changes in environment outside of the typical range a cell is exposed to. We propose phenotypic pliancy as a clarifying distinction between these phenomena, although other papers may refer to both as phenotypic plasticity.

While much knowledge has been gained about the mechanisms and functions of the various PRCs and their PcG protein components, much still remains unknown owing to the complexity of the polycomb system. For humans and other complex multicellular organisms, the set of all possible combinations of PcG proteins that can form different PRCs and how they function is still largely unknown (there are estimated to be more than 100 different PRC variants) [3, 14, 17, 18]. Research is ongoing for ways of determining PRCs’ target gene sets, which are thought to be different but possibly overlapping between the various PRCs [18, 19]. Additionally, despite much research, there is no determined DNA sequence that distinguishes a target gene (called Polycomb Repressive Elements (PREs)) like in *Drosophila Melanogaster* [20–22] and no current way to reliably classify PRC target genes in humans [18, 20, 23].

Given the lack of target gene knowledge and the immense number of possible combinations of PcG proteins combining to form different PRCs with potentially differing functions and target gene sets in different contexts (developmental stages, cell types, and environments), it is difficult to experimentally study in a systematic way a breakdown of the polycomb mechanisms and its consequences on phenotypic fidelity in humans. As an alternative, we introduce a new conceptual computational model that allows us to test our hypothesis *in-silico* with full control over the model’s parameters. By using a computational model, we can investigate the *general* functions of polycomb mechanisms, and emergent properties upon their evolution and breakdown that are independent of specific contexts. Our model aims to capture the essential relevant aspects of epigenetic polycomb mechanism control in differentiation, fidelity, and pliancy without being encumbered by precise biological details under specific conditions. Additionally, our model aims to produce robust qualitative results, rather than quantitatively precise predictions.

One possible occurrence of phenotypic pliancy is in metastatic cancer. In contrast to the phenotypic fidelity exhibited by normally functioning cells, metastatic cancer cells seemingly exhibit phenotypic pliancy [24–29]. The wide variety of environments encountered is highlighted throughout the course of the disease as the metastatic cells progress through invasion, intravasation, survival in the circulatory system, extravasation, arrival at the metastatic site, and, finally, colonization of the metastatic site [30–32]. Each of these occur in a different environment and seemingly requires the metastatic cells to assume a different phenotype in order to adapt to and survive in that environment [24–29, 31], while non-metastatic cancer cells do not switch their phenotype in response to environmental change [31]. Additionally, it has been observed that sustained abnormal phenotypic switching is not limited to cancer stem cells but is the property of a wider cell population [24, 28, 29]. Attaining a deeper understanding of the underlying mechanism of such abnormal phenotypic pliancy is therefore paramount.

As a special case of our general hypothesis, we hypothesize that progression to metastasis is driven by the emergence of phenotypic pliancy in cancer cells as a result of polycomb mechanism dysregulation.

Indeed, current literature demonstrates that dysregulation of polycomb mechanisms plays an important role in cancer and metastasis [3, 9, 14, 15, 17, 18, 33], but explanations of how and why remain contradictory. A potential explanation of the discrepancies in polycomb mechanism involvement in primary and metastatic cancers, is that PcG proteins, when not involved in a PRC, have different functions when acting in isolation versus as part of a PRC [17]. For example, EZH2 is a PcG protein component of the Polycomb Repressive Complex 2 (PRC2) that on its own is thought to be involved in cell cycle progression [4]. So EZH2 is often up regulated in primary tumor cancers, which may help cancer cells proliferate [34–36]. However, the other PRC2 PcG protein components are not up regulated, hinting that EZH2 up regulation may not be affecting the PRC2 function. When EZH2 is found to be down regulated, it is typically in conjunction with the other PRC2 components and in metastatic cancers [4].

Previous hypotheses of disease progression behaviors that could possibly explain metastatic cancer cells’ phenotypic pliancy are de- and re-differentiation [37–39]; cells’ accumulation of appropriate mutations while dormant [40, 41]; or exosome-mediated metastasis [42, 43]. However, there are important commonalities captured by our hypothesis that are involved in these three seemingly different behaviors: environmental influences, epigenetic modifications, and abnormal phenotypic switching. Our *systems-level* hypothesis does not contradict these more circumscribed hypotheses, but rather may provide a more parsimonious and unifying conceptual mechanistic framework that may shed light on the mechanisms underlying all the above previous hypotheses.

In-silico experiments using our computational model demonstrate, in a general and context-independent way, that the breakage of polycomb mechanisms leads to the emergence of phenotypic pliancy. As hypothesized, the observed pliancy is sustainable, with phenotypic movement towards the evolved normal phenotype in the surrounding environment, and the degree of phenotypic movement is correlated with the extent of polycomb dysregulation.

We then corroborate our in-silico results by examining solid tumor metastatic cancers, where we hypothesize phenotypic pliancy is implicated in metastasis. Since phenotypic pliancy is a cellular-level property, we use single-cell RNA-sequencing data. Our analysis demonstrates that metastatic cancer cells display phenotypic pliancy behavior correlated with the breakage of polycomb mechanisms.

## Results

### Model Description

We develop a holistic computational model of the developmental process including the action of PcG-like mechanisms (referred to as PcG mechanisms for the rest of model description and results), as well as post-developmental gene-regulatory dynamics, and the evolution of these dynamics for populations of individuals. A conceptual summary of our model follows (see Materials and Methods computational model description section for the mathematical detail).

A cell is modeled by a gene regulatory network, represented by a gene-by-gene matrix whose entries define the influence of the product of gene j on the expression of gene i. This network is augmented by a gene-environment interaction matrix that models how the expression of gene i is influenced by an environmental state vector that is constant in time (see Figure 2, Founder in Env. 1 & 2).

**Fig 2.**
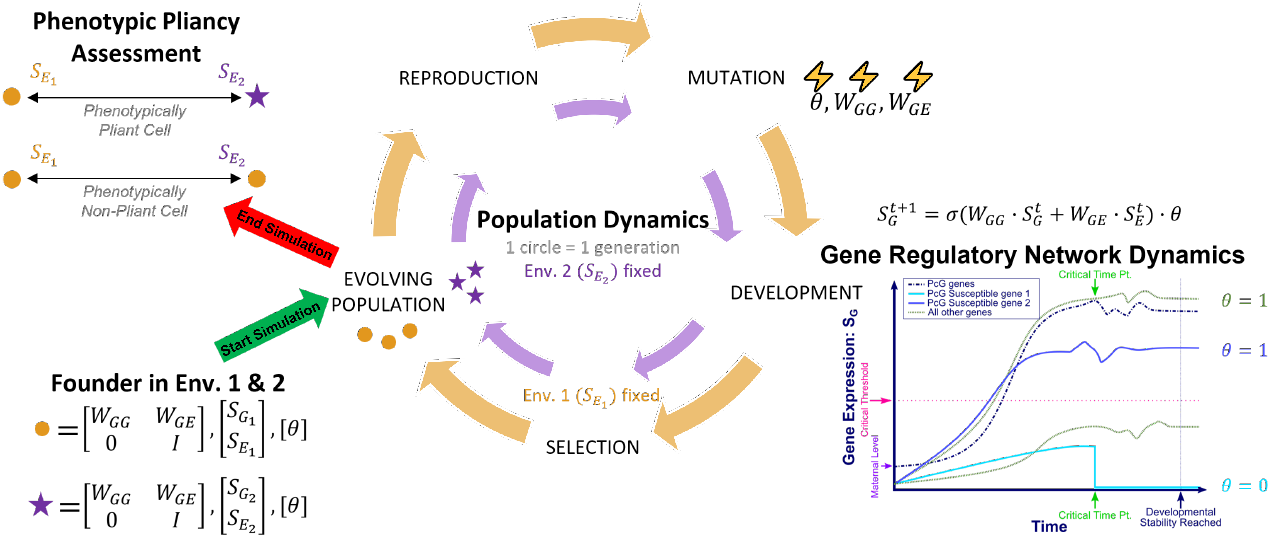
Schematic of our Computational Model: Schematic of of our computational model, where each simulated cell’s phenotype in the population is represented by an orange circle for environment 1 and a purple star for environment 2. Evolution starts with a single founder that has two different “cell types” that differentiate in environment 1 and environment 2. The population dynamics portion of the schematic represents the steps in each generation of evolution. The gene regulatory network dynamics portion of the schematic depicts development of a single individual in the population and shows how PcG mechanisms can alter the resulting gene expression based on if a PcG target is repressed during development or not (repressed if *θ* is zero and free of PcG control if one). This figure also shows a simplified schematic of phenotypic pliancy assessment when a simulated cell switches from environment 1 to environment 2. To generate our in-silico data, we simulate 10,000 evolved populations, by starting with 1,000 different initial populations with each undergoing 10 different evolutionary trajectories. Each population consists of 1,000 individuals and undergoes 1,000 generations. Please see the Model Description section for a conceptual description of the model and the Materials and Methods section for a mathematical description of the model.

The expression of each gene also depends on whether or not it is controlled by a PcG mechanism complex that makes it susceptible to epigenetic regulation. Given an initial gene expression state vector and environmental state vector, the model gives rise to a deterministic dynamical process, which is thought of as “development” (see Figure 2, Gene Regulatory Network Dynamics). The developmental process is given by dynamically updating the gene expression vector each time step, summing up all influences (genetic and environmental) on each given gene and mapping the result through a sigmoidal function to determine the final effect on that gene. The epigenetic regulation, if any, of a gene, comes into effect after a certain developmental time has elapsed. If at that time point a gene is expressed below a certain threshold and is controlled by a PcG mechanism complex, it is permanently silenced, effectively punching a hole into the network (see Figure 2, Gene Regulatory Network Dynamics). During development, the environment also influences the expression of the genes and therefore which genes become permanently silenced. This dynamical process eventually settles into a stable steady-state gene expression vector, which constitutes the (high-dimensional) “phenotype” of a cell.

Together, all gene regulatory network matrices, including the matrix governing environmental susceptibility and the matrix establishing the epigenetic footprint, constitute the “genotype” of a cell (see Figure 2, Founder in Env. 1 & 2). The dynamical process governed by the “genotype” within a fixed environment constitutes “development”, which results in a “phenotype” (see Figure 2, Gene Regulatory Network Dynamics). This scheme therefore establishes a holistic genotype-phenotype map.

We then define an “individual” as consisting of two cells – call them cell 1 and cell 2 – with identical genotype, as a primitive model of multicellularity. A population of such (initially identical) individuals is generated (see Figure 2, Population Dynamics). Two environments are defined and for each environment we specify an optimal target gene expression state vector (i.e. a desired phenotype). This setup is meant to capture the idea that within each individual, cell 1 develops in environment 1 (generating phenotype 1) and cell 2 in environment 2 (generating phenotype 2). One might think of these as two identical stem cells which differentiate into distinct phenotypes according to which of the two environments they develop in. The population is now evolved by mutating the genotype which is shared by cells 1 and 2 within each individual (see Figure 2, Population Dynamics, Mutation). This evolutionary process aims to attain a genotype such that each cell of an individual attains as closely as possible the target phenotype appropriate for the environment it is exposed to during development. This is meant to represent the evolution of differentiated multicellularity.

Our model is built upon a well-established computational gene regulatory network model. The original model was used to study the evolution of canalization [44]. Later, it was used to study the role of a PRC in decoupling genetic and environmental robustness during development [16]. The main additional features of our current, extended computational model are: explicit environment-gene interations, multiple PcG epigenetic mechanism complexes, and multicellularity.

In total, these features allow us to carry out in-silico multicellular evolution experiments with evolved PcG epigenetic regulation to investigate phenotypic fidelity and phenotypic pliancy when the PcG mechanism is intact and broken (see Figure 2, Phenotypic Pliancy Assessment), as described in the next section. Given that much remains unknown about PcG mechanism complexes and their targets in humans leading to incomplete gene regulatory networks, we build and leverage a computational model in order to investigate phenotypic fidelity and pliancy from a generalized perspective over a large set of parameters and network architectures. Importantly, the knowledge we gain will not be contigent on specific networks, environments, disease conditions, experimental conditions, or organism.

### Emergence of Phenotypic Pliancy during Evolution: Model Results

We investigate the role of PcG mechanisms in the evolution towards phenotypic fidelity to differentiated phenotypes, and the general consequences of dysregulating these evolved PcG mechanisms post-developmentally. We hypothesize that post-developmental dysregulation of PcG mechanisms allows cells to become phenotypically pliant. We use our computational model described above to generate artificial “single-cell gene expression” data. With this generated data, we visually and quantitatively demonstrate that intact PcG mechanisms ensure phenotypic fidelity behavior, while PcG mechanism dysregulation causes phenotypic pliancy to emerge.

Using our computational model, we simulate 10,000 evolved populations, by starting with 1,000 different initial populations and letting each of them evolve under 10 different evolutionary trajectories, each determined by different random seeds which result in differences in mutation and reproduction over the course of evolution. Each population consists of 1,000 individuals and undergoes 1,000 generations of evolution, i.e. 1,000 cycles of the population dynamics given by our model and depicted schematically in Figure 2.

Throughout the course of evolution, the total population fitness results indeed tend to increase as expected (see Supplementary Figure S1). Increasing fitness means that the populations successfully evolve toward the selected-for differentiated phenotypes. Note, the fitness in environment 1 is expected to decrease since all individuals start at the optimum in environment 1 but not in environment 2, so it must evolve towards environment 2, i.e. demonstrating the evolution towards the capacity for differentiation (see Materials and Methods for details).

Moreover, we see that during the course of evolution, PcG mechanisms take an increasingly active role, with an increasing number of genes under PcG mechanism control (see Supplementary Figure S2), and correspondingly a decreasing effective connectivity of the gene-regulatory network (see Supplementary Figure S3). The effective connectivity is the result of removing the genes which are repressed by PRCs during development in each environment and their connections in the network, similar to the representation in Figure 1. Interestingly, our model predicts that there is a preference for a higher incoming connectivity among PcG mechanism target genes that are repressed by any PRC, relative to non-repressed target and non-target genes (see Supplementary Figure S3, A). A possible explanation for a higher incoming connectivity of genes repressed by PcG mechanisms could be that these repressed genes are acting as switches in the gene regulatory network to turn on and off a particular downstream pathway based on the environmental context. So the average incoming connectivity increases for these genes to facilitate two different differentiated expression profiles based on the same gene regulatory network under two different environments. These trends support previous biological and computational observations that PcG mechanisms play an important role in facilitating the evolved differentiation.

Next, we observe that the evolution of populations with evolved PcG mechanisms promotes phenotypic fidelity as an emergent property, without being directly selected for. To see this, we apply principal component analysis (PCA) to the gene expression vectors of cells in a representative population. We carry out this analysis at two evolutionary time points, after 50 and 1,000 generations, respectively. After 1,000 generations (see Figure 3A), we observe phenotypic fidelity. Specifically, when PcG mechanisms are left intact post-developmentally and the cells which developed in environment 1 are transferred to environment 2, the resulting stable gene expression patterns (cyan) largely remain close to the original gene expression patterns of cells developed in environment 1 (orange) while largely staying far from the gene expression patterns of cells developed in environment 2 (purple X’s).

**Fig 3.**
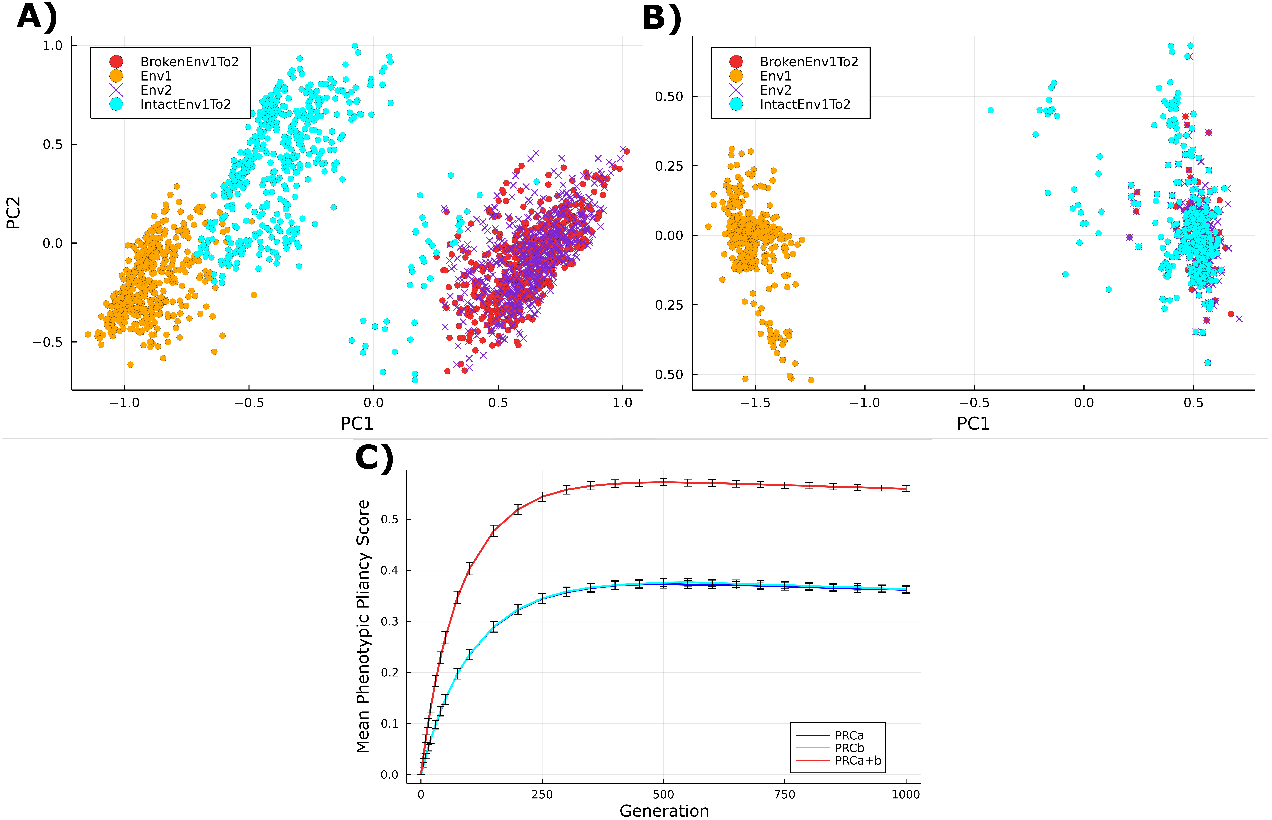
Phenotypic Pliancy and Fidelity Results from Computational Model: **A**. PCA results for all individuals in population at the end of evolution (after generation 1,000) when polycomb mechanisms have more fully evolved to visually assess phenotypic pliancy. When transfer from environment 1 to environment 2, only the case when polycomb mechanisms are broken and switched to environment 2 (red) alter their phenotypes to resemble that of environment 2 that moved to (purple X’s). The individuals with polycomb mechanisms left intact and moved to environment 2 (cyan) have phenotypes that do not switch and resemble more closely environment 1 that moved from (orange). **B**. Principle Componenet Analysis (PCA) results for all individuals in population right after generation 50 during evolution when polycomb mechanisms have not more fully evolved to visually assess phenotypic pliancy. When transfer from environment 1 to environment 2, both the cases when polycomb mechanisms are left intact (cyan) and broken (red) alter their phenotypes to resemble that of environment 2 that moved to (purple X’s). **C**. Average phenotypic pliancy score when vary degree of PcG-like mechanisms dysregulation during evolution for all 10,000 populations when move from environment 1 to environment 2 to quantitatively assess pliancy. We vary degree of dysregulation by breaking PRCa alone (blue), PRCb alone (cyan), or both PRCa and b (red) for all 10,000 populations for a total of 10 million simulated cells pliancy score averaged (y-axis) for different generations throughout evolution (x-axis).

In contrast, the population at an early stage of evolution (see Figure 3B) does not exhibit phenotypic fidelity because the polycomb mechanism has not yet evolved. When the same experiment is carried out, the cells which developed in environment 1 and transferred to environment 2 with PcG mechanisms intact adopt gene expression patterns much closer to those of cells developed in environment 2 (purple X’s). Similarly, we ran a control experiment in which the evolutionary dynamics are carried out in the absence of any PcG mechanism evolution, and we see the same results of no phenotypic fidelity even after 1,000 generations (see Supplementary Figure S4).

These results confirm biological understanding of the role of an evolved polycomb mechanism in maintaining cell identity, and shows that our computational model, while general and abstract, is sufficiently true to biological reality to manifest this emergent property.

Finally, we observe that, when evolved populations have PcG mechanisms broken post-development, they exhibit phenotypic pliancy. This can be seen both visually in a PCA, as well as numerically (see Figure 3A and 3C). In the PCA, when PcG mechanisms are broken post-developmentally and the population is then transferred from environment 1 to environment 2, the resulting stable gene expression patterns (red) move closer to the normal phenotype in environment 2 (purple X’s), and away from their original phenotype in environment 1 (orange). This observation illustrates that, when a PcG mechanism is broken and the population is transferred to a different environment, cells will switch their phenotypes to phenotypes that more closely resemble the normal phenotypes that evolved in the environment transferred to.

For numerical evidence, we devise a measure of pliancy that measures, for each individual, the proportion of genes which, when PcG mechanisms are broken post-development and upon transferring to a different environment, move away from the individual’s phenotype in the original environment and toward its phenotype in the new environment and more so than when PcG is left intact (see Supplementary Figure S5 and Materials and Methods). We see that this phenotypic pliancy score tends to increase substantially over the course of evolution, indicating that phenotypic pliancy upon broken PcG mechanisms is an emergent property of evolution (see Figure 3C). Specifically, we compute a phenotypic pliancy score upon breaking both PcG mechanisms, and when breaking each PRC separately (see Materials and Methods). We find that the level of phenotypic pliancy corresponds to the degree of polycomb mechanisms dysregulation, as the average phenotypic pliancy of all 10 million simulated individuals is much greater when both PRCa and PRCb are dysregulated simultaneously than when PRCa or PRCb is dysregulated separately (see Figure 3C). The dysregulation of these PRCs does not affect the stability of the simulated cells, where around 0 to 0.1% of cells are unstable upon dysregulation of the PRCs separately or together (see Supplementary Figure S6).

Importantly, we find that phenotypic pliancy is sustained, meaning that cells can keep switching their phenotype when placed in a different environment and not get stuck in a certain phenotype. This sustainability can be visualized using PCA for the case when switch back to environment 1 after switching to environment 2 (see Supplementary Figure S7), such that the phenotype switches back to resemble that of environment 1 (orange circles) when moved back to environment 1 (blue X’s). Lastly, as a control, we test the case when PcG mechanisms are broken but the cells are kept in the same environment in which they had developed (green circles). We find the gene expression patterns do move away from their normal expression, but not substantially (see tan circles in Supplementary Figure S7).

To verify our model results are robust, we test for phenotypic pliancy over a wide range of parameters, demonstrating that phenotypic pliancy is a general, systems-level phenomenon (see Supplementary Figure S8 and Supplementary Information).

In sum, our model results demonstrate that: 1) evolution of PcG mechanisms causes post-developmental phenotypic fidelity to evolve, 2) a breakdown of PcG mechanisms lead to the emergence of phenotypic pliancy, 3) the level of phenotypic pliancy corresponds to the degree breakdown of PcG mechanisms, 4) phenotypic pliancy is sustainable, and 5) not only do phenotypically pliant cells move away from their evolved phenotype in the environment they move from, but they also move closer to the normal evolved phenotype in their new environment.

Importantly, our model appears to capture some biological phenomena of differentiation, phenotypic fidelity, and phenotypic pliancy, and it does so in a robust way.

### Assessment of Phenotypic Pliancy in Metastatic Cancer

To assess our hypothesis and our model’s predictions, we use publicly available single-cell RNA-sequencing data from matching primary tumor and metastatic cancer sites, and normal cells at each site. This data departs from our in-silico data in various ways. In this data we do not have access to identical cells placed in different environments. Instead, we analyze populations of individual cells and make use of the fact that metastatic cells have undergone a change of environment from the primary to the metastatic site. Moreover, unlike with our model data, where the only experimental conditions involve polycomb dysregulation and a change in environment, here we are faced with additional mutations and abnormalities coming from cancer cells. To control for that, we compare metastatic cells both to the primary tumor cells and to the normal cells at both the primary and metastatic sites. Driven by our findings from the model in the above results section, we address the following three sub-hypotheses: 1) PcG genes are differentially expressed in cancer cells relative to normal cells in their environment; 2) metastatic cells exhibit elements of phenotypic pliancy, such that their phenotype (i.e. gene expression profile) moves away from that of the primary tumor, and closer to the normal cells in the metastatic site; and 3) PcG genes are implicated in the observed phenotypic pliancy of metastatic cells, such that there is a positive correlation between the extent of dysregulation of PcG genes and the degree of phenotypic movement in the direction of normal phenotype.

Here we emphasize that metastatic cells’ phenotypic movement in the direction of normal does not imply taking on the identity of the surrounding normal cells.

We use two different single-cell RNA-sequencing data sets from patients with metastatic head and neck cancer (H&N) obtained by Puram et. al. [45] and patients with metastatic serous epithelial ovarian cancer (ovarian) obtained by Shih et. al [46] (see Materials and Methods). We utilize a list we curated with 69 genes that are involved in polycomb mechanisms including 22 PcG genes, 35 TrxG genes, and 12 genes that have been verified to be controlled by PRCs (see Supplementary Table 1) [14, 15, 19, 47]. Preprocessing and all further analysis of the Puram et. al. and Shin et. al. data sets is performed using the R software package Seurat (see Materials and Methods) [48–50].

#### Metastatic cells are enriched with dysregulated PcG mechanisms

We test sub-hypothesis 1 by performing differential expression analysis on the preprocessed data using the default method (Wilcoxon rank sum test) of the R Seurat package [48–50]. For the H&N data set, differential gene expression analysis is done separately for each of the two pairwise comparisons: metastatic cells versus non-cancer cells in the lymph nodes, and primary tumor cells versus non-cancer cells in the oral cavity. For the ovarian data set, differential gene expression analysis is done between primary tumor cells versus normal cells in the ovary, and metastatic cells versus these same normal cells in the ovary since normal samples were not taken in the omentum.

For both H&N and ovarian data, our results for primary versus normal cells (column 2, row 1 and 3 in Supplementary Table 2, respectively) and metastatic versus normal cells (column 2, row 2 and 4 in Supplementary Table 2) show no significant trend in the number of polycomb mechanism genes that are differentially expressed. Since down regulation is more indicative of a breakdown in function, we investigate the extent to which these differentially expressed genes are up regulated or down regulated. To do this, we calculate the log-fold change sum for primary and normal cells, separately, at the primary site (column 3, row 1 and 3 in Supplementary Table 2) and the sum for metastatic and normal cells at the metastatic and primary site, respectively (column 3, row 2 and 4 in Supplementary Table 2). To calculate the significance in the difference in the log-fold change sums between primary and metastatic for each H&N and ovarian, we test against the null hypothesis that polycomb dysregulation is the same in primary and metastatic, by using a permutation test randomly permuting, over 1000 trials, the identity of primary and metastatic for each PcG gene’s expression data, and using the generated histogram to extract p-values.

For the H&N results, a decrease in log-fold change sum at the metastatic site (p-value *<* 0.15) gives partial evidence that metastatic cells’ PcG mechanism is being down regulated to a greater extent than primary cells. For the ovarian results, despite a decrease in the number of differentially expressed PcG genes in metastatic cells compared to normal, we observe strong evidence that metastatic cells’ PcG mechanism is being highly down regulated while primary cells’ mechanism is actually being up regulated (p-value *<* 0.05). Since down regulation is more indicative of breakdown in function, this supports our hypothesis of increased polycomb dysregulation in metastasis. Up regulation in primary cells may be explained by many findings that some polycomb mechanism genes have independent functions from their PRC(s) in many cancers, such as cell cycle progression roles, as discussed for EZH2 [4, 34–36].

#### Metastatic cells are phenotypically pliant

To test sub-hypothesis 2 and assess our model prediction that cells with broken polycomb mechanisms move toward the normal phenotype of the new environment in which they are placed (see Figure 3A and B), we further narrow the set of genes under consideration to only those which are differentially expressed between primary and metastatic cancer cells resulting in a total of 2,246 genes for the H&N and 3,393 genes for the ovarian data. This restriction is in order to not drown out the signal of the difference between these two cellular categories.

In order to visualize phenotypic pliancy, we use 2 different methods to allow us to capture critical features of the data to extract knowledge of phenotypic pliancy behavior without having dynamical data. First, we perform PCA on the two data sets. The first two principal components (PCs) are plotted in Supplementary Figure S9, showing a clear clustering of the data for both the H&N and ovarian data. Importantly, along the first PC for the H&N data, the majority of the metastatic cells tend to be closer to normal cells than primary tumor cells are Supplementary Figure S9, A. While, in the ovarian data there is little difference between metastatic, primary, and normal in the first PC but in the second PC the normal cells overlap with the metastatic cells Supplementary Figure S9, B. These PCA results lend credence to our hypothesis of increased phenotypic pliancy in metastatic cells and prediction from our model. Moreover, Supplementary Figure S9 reveals that primary tumor cells can be divided into two sub-clusters based on the sign of PC2 (y-axis), where one sub-cluster substantially overlaps with metastatic cells (in cyan).

Secondly, we apply Uniform Manifold Approximation and Projection (UMAP) [51] to the two data sets. The UMAP algorithm, as opposed to PCA, preserves more of the local structure of the data rather than just the global linear structure, and typically shows a continuous representation of the local structure, mapping nearby points to nearby connected points. Since the primary, metastatic and normal cells are arranged in a connected cluster in the UMAP results for both H&N and ovarian data, we can compare these three categories to each other and the results reveal a clear continuous progression from primary tumor cells to metastatic cells and then to normal cells (see Figure 4). These UMAP results give further credence to metastatic cells being closer to normal cells than primary tumor cells, and thus our hypothesis that metastatic cells have increased phenotypic pliancy. In addition, these UMAP findings also reveal that a significant number of primary tumor cells (labeled cyan in Supplementary Figure S9) possess a gene-expression profile closer to that of metastatic, which suggests these primary cells may have transitioned to a more metastatic cell type (see Figure 4).

**Fig 4.**
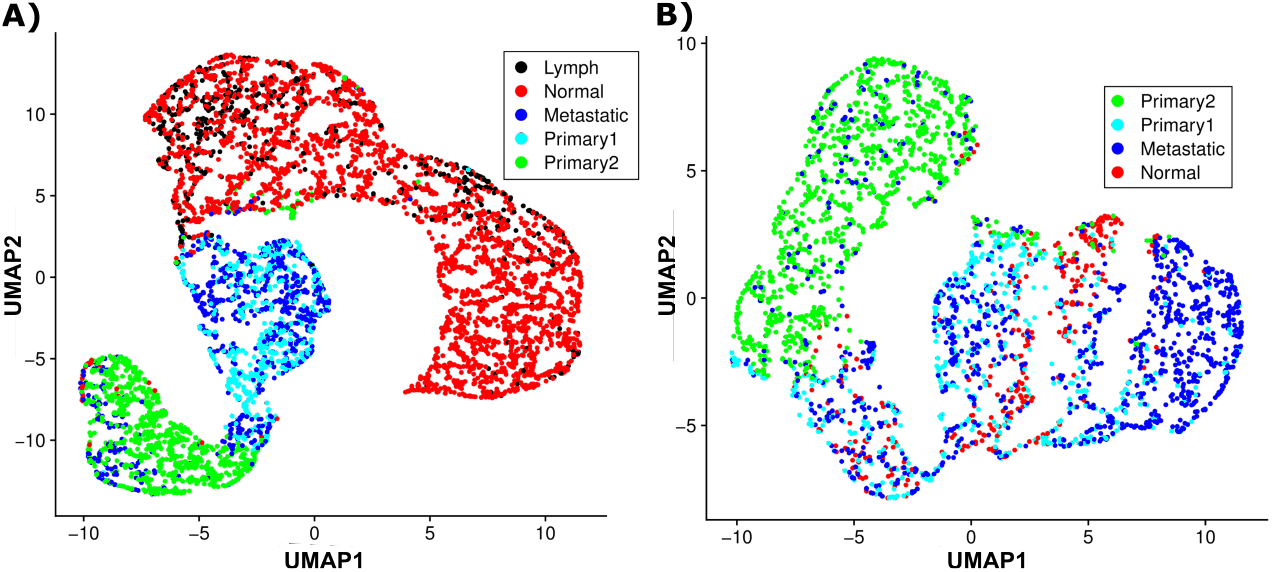
UMAP Results for Metastatic Cancer SC-RNA-Seq Datasets: UMAP results for the two single-cell RNA-sequencing data sets: **A**. H&N data and **B**. ovarian data. Note: the primary tumor cells are colored based on the PCA sub-clustering in Supplementary Figure S9.

We quantify if metastatic cells are phenotypically pliant using two methods. Firstly, we measure whether the distance of metastatic phenotype (gene expression profile) to the normal phenotype is less than the distance of primary phenotype to normal phenotype. We compute the distance between two populations of cells as the Euclidean distance between the centroids of their log-scale gene expression data. With this measure, the distance between primary tumor and normal cells is 55.1, which is larger than the distance between metastatic and lymph normal cells, which is 50.2 for the H&N data. For the ovarian data, the distance between primary tumor and normal cells is 34.2, which is larger than the distance between metastatic and normal ovary cells of 32.5. These distances are further increased when only considering the “primary2” cells that are clustered separately from the metastatic cells (green in Supplementary Figure S9), with “primary2” to normal distance of 57.4 compared to metastatic lymph normal distance 50.2 for H&N, and primary2 to normal distance of 39.8 compared to metastatic normal distance 32.5 for ovarian data.

Secondly, we calculate pliancy at the gene level, such that a gene is pliant if the distance of its expression in metastatic cells to normal cells is less than the distance of its expression in primary cells to normal cells. We calculate the gene-by-gene differences between primary and normal at the primary site (x-axis) versus the differences between metastatic and normal at the lymph node site or ovary site for H&N and ovarian data, respectively (y-axis) (see Supplementary Figure S10), and find, of the genes differentially expressed between primary tumor versus metastatic cells, 68% are pliant (p-value *<* 10^−8^) for H&N and 72% (p-value *<* 10^−9^) for ovarian. All these measures lend strong evidence to sub-hypothesis 2 of increased phenotypic pliancy of metastatic cells for two different cancer types and our model predictions.

#### Degree of PcG mechanism dysregulation correlates with level phenotypic pliancy

To test our model prediction that the degree of phenotypic pliancy is correlated with the level of polycomb mechanisms disruption (see Figure 3C), we assign a measure of phenotypic pliancy to each metastatic cell, and test the correlation of this measure to the level of PcG mechanism dysregulation.

We measure a metastatic cell’s phenotypic pliancy as the *pliancy z-score*, that measures the relative extent of movement of a particular metastatic cell’s phenotype toward the normal phenotype and away from the mean of all metastatic cells (see Materials and Methods for details). We use the genes differentially expressed between metastatic and normal cells, excluding the 53 PcG and TrxG genes because we use these PcG mechanism genes as predictors in the regression analysis below (these 53 genes do not include the 12 known PcG target genes) for the H&N data and excluding the 56 PcG and TrxG genes for the ovarian data (these do not include the 9 known PcG target genes). We then fit a linear regression to predict the pliancy *z*-score as a function of all the PcG mechanism genes’ expressions (*r*^2^ = 0.22 for H&N and *r*^2^ = 0.56 for ovarian data). For the H&N data, out of the 53 PcG and TrxG genes, 30 have positive coefficients in the linear model. Positive coefficients mean that decreased PcG and TrxG expression is positively correlated with metastatic cells’ lower distance to normal cells, thus indicating that dysregulation of the PcG mechanism drives pliancy. For the ovarian data, out of the 56 PcG and TrxG genes, 21 have positive coefficients.

As a control for this result, we perform a bootstrap, randomly picking 1000 samples of 53 genes for H&N and 56 genes for ovarian and performing the same linear regression of the score as a function of these 53 and 56 genes’ expression profiles for H&N and ovarian data, respectively. For each such regression, we record the number of genes with a positive coefficient and compute the distribution of this number (see Supplementary Figure S11). The mean is 22.3 out of 53 for H&N data, and mean is 10.6 out of 56 for the ovarian data. Relative to these distributions, 30 for H&N data and 21 for ovarian data is statistically significant (p-value *<* 0.05 and p-value *<* 10^−5^, respectively) (see Supplementary Figure S11). So, in both the H&N and ovarian data there is a high level of dysregulation of PcG mechanism genes that is associated with metastatic cells moving closer toward the normal phenotype.

## Discussion

In this paper, we propose and validate a novel framework of phenotypic pliancy, and its emergence as a result of PcG mechanism dysregulation.

We introduce an abstract computational model of the evolution of gene-regulatory networks together with PcG mechanisms. Using this model, we demonstrate that the evolution of PcG mechanisms results in phenotypic fidelity to differentiated phenotypes, while dysregulation of PcG mechanisms leads to decreased phenotypic fidelity, which allows dysregulated cells to become phenotypically pliant. Additionally, we show that cells with dysregulated PcG mechanisms, when introduced into a new environment, switch into phenotypes that more closely resemble that of the already evolved phenotypes in this environment, which we expect since the environmental cues should dominate once the cellular memory disappears due to loss of polycomb mechanisms.

Our model results demonstrate that phenotypic fidelity and phenotypic pliancy are both general emergent properties over a wide range of simulated gene-regulatory networks and polycomb mechanism patterns, suggesting that these are general systems-level phenomena independent of specific contexts or selection. Similarly, previous research using abstract biologically inspired models demonstrated the emergence of systems-level properties that resulted in important biological insights [16, 44, 52–54].

We begin to test if the model conclusions hold true in biological contexts by utilizing publicly available single-cell RNA-sequencing data from metastatic cancer. We find preliminary evidence that 1) metastatic cells have enrichment of PcG mechanism genes differentially expressed when compared to normal, non-cancer cells at the respective site relative to primary tumor cells; 2) metastatic cells behave like phenotypically pliant cells, such that their phenotypes move away from that of the primary tumor, and closer to the normal, non-cancer cells in their surrounding site; and 3) PcG mechanism dysregulation is positively correlated with the degree of phenotypic movement in the direction of normal phenotype, i.e. level of phenotypic pliancy.

The metastatic cancer data findings corroborate the results from our computational model. While our metastatic cancer data results are not definitive since many PRCs, PcG mechanism genes, and PcG target genes are still unknown, our results do provide initial evidence of a general hypothesis that can be further tested and validated with experimental studies. If correct, these findings can explain a mechanism driving development and progression of metastatic cancer. Additionally, our computational model can be used in future studies to help guide the criteria for identifying polycomb mechanism target genes that contribute most to phenotypic pliancy. For example, our model predicts that these target genes will have higher incoming connections relative to the rest of the genes in the gene regulatory network under investigation.

Our findings related to metastatic cancer cells being phenotypically pliant do not contradict but rather may provide a more parsimonious and unifying conceptual mechanistic framework for metastatic disease emergence and progression than the existing more circumscribed hypotheses, i.e. de- and re-differentiation; appropriate mutation accumulation during dormancy; and exosome-mediated niche-construction. Future work can investigate the interplay between polycomb mechanisms dysregulation and the above hypotheses in driving phenotypic pliancy and metastatic cancer.

There are additional unknowns pertaining to the primary tumor that need to be addressed in future work to understand the initial stages of metastatic cancer development. Further research is needed to assess whether polycomb mechanisms evolve to provide phenotypic fidelity to the generated tumor microenvironment of primary tumor cells. Additionally, our computational model does not address the interplay between polycomb mechanisms dysregulation and the accumulated mutations that lead to development of primary cancer. Thus, further detailed analysis is needed to investigate the implications of this interplay.

We demonstrate that the emergence of phenotypic pliancy is not contingent on selection pressures, contexts, or levels of investigation and therefore could hold true in diverse and broad scopes. Thus, our proposed mechanism of phenotypic pliancy has the capacity to be a general phenomenon not unique to cancer metastasis, and may shed light on complex diseases where sustained abnormal phenotypic switching has been observed [55]. Even though our results are reproducible in both our computational model and two different solid tumor metastatic cancers, further experimental work and testing is needed to assess phenotypic pliancy’s broader implications.

## Materials and methods

### Computational Model Description

Our computational model operates on two levels: gene regulatory network (cell) dynamics and population dynamics (see Figure 2). At the detailed level of gene-regulatory network dynamics, a simulated cell’s state at any given time is given by a gene expression state vector, S_W_, together with an environment state vector, S_E_. The gene expression vector has dimension *N* – the number of genes – and its coefficients represent gene expression levels. The environment vector’s coefficients represent abstract “environmental factors” which can potentially affect gene expression. The dimensionality of these environmental factors is a parameter, which we choose to be equal to *N*. Next, the basic gene-regulatory dynamics are described by two matrices (see Figure 2). The first matrix, *W*_*GG*_, is a *N × N* matrix encoding interactions among genes, where the *ij* entry of the *W*_*GG*_ matrix represents the effect of the product of gene *j* on gene *i*. The second matrix, denoted *W*_*GE*_, similarly encodes the effect of the environmental factors on gene products in the cell. Please see Supplementary Table 3 for a list of model parameters and their values.

Finally, we incorporate the effects of PcG proteins in a developing individual into the model. Each simulated “cell” in this model is assigned multiple simulated PRC’s. The information of which genes are targets of a given PRC is encoded in a PRE matrix (denoted *θ*), with entry *θ*_i,j_ equal to 1 if gene *i* is under the control of PRC *j*, and equal to 0 otherwise. During the developmental step in our model, a gene possessing PREs (*θ*_i,j_ = 1 for some *j*) is permanently silenced by a PRC if that gene fails to be expressed above a threshold *γ* by a predetermined critical time point t_c_ during development.

Letting 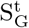 denote the gene-expression state vector at time *t* and assuming a fixed environment state vector S_E_, the gene expression dynamics of each simulated cell are then given by 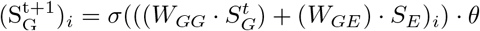, where *σ* is the standard sigmoid. If gene *i* is under the control of a PcG mechanism, *t > t*_*c*_, and 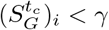, then 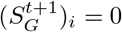.

Development is defined as the process of starting from some initial state and iterating the gene expression dynamics as described above until steady state is reached, denoted *S*_*G*_. Steady state is reached when a stability measurement, a normalized variance of gene-expression pattern within the last 10 developmental time-steps, is smaller than the error term *ϵ* = 10^−4^. Cells that reach steady state are deemed developmentally stable [52, 56], otherwise they are considered lethal.

At the detailed level of population dynamics, a population of *M* individuals undergo iterations of mutation, reproduction, development, and selection, where each iteration represents a generation (see Figure 2). In total, the population dynamics step models the evolution of the population through 1,000 generations in the presence of selection. A population is initially generated from one developmentally stable individual, known as a founder. The matrices *W*_*GG*_ and *W*_*GE*_, together with the PRE matrix *θ*, constitute the individual’s genotype. The *W*_*GG*_ and *W*_*GE*_ matrices of the founder are randomly generated matrices with a fixed ratio *c* (the matrix connectivity) of non-zero, Gaussian-distributed entries. The matrix *θ* is initially set to 0, with a fixed number of potential PRCs, so that in the beginning of evolution no gene is under PcG mechanism control. We add the ability to model the evolution of a population of multicellular individuals (see Figure 2), each individual’s cells sharing the same genotype but developing in two different environments. We randomly generate 2 different environment binary state vectors which are pairwise different by a certain percentage difference, Δ_*e*_, and such that the founder individual is stable in these environments. We set the optimal phenotype in environment 1 to be equal to the stable state of the founder in environment 1, and we randomly pick the optimal phenotypic vector in environment 2, 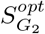, such that it is different by a threshold, Δ_*S*_, from environment 1. We pick initial state vectors 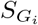 which are environmentally stable in their respective environments but may differ from 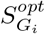, allowing for evolution toward the optimum.

We measure developmentally stable individuals’ fitness Ω with stabilizing and directional selection components:

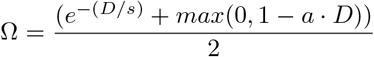

where 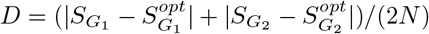 measures (*L*^1^) distance from optimum, *s* is the stabilizing selection strength, and *a* is a parameter of the directional selection strength. Note here *D* simply measures the total deviation between the observed phenotypes and optimal phenotypes in both environments, normalized by the number of genes. The fitness term *e*^−(*D/s*)^ represents stabilizing selection since it drops off *exponentially* if *D* deviates from 0. On the other hand, the fitness term *max*(0, 1 − *a* · *D*) represents directional selection, as it rewards moving towards the optimum in a *linear* way as long as *D* is not too large 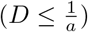. Thus the directional selection term helps direct a population which is far away from optimum towards optimum, while the stabilizing selection term helps more strongly maintain an optimum phenotype once it is reached. These selection pressures apply in both environments at once. Note in our case there is an asymmetry: as environment 1 starts at optimum, stabilizing selection is the relevant force, while environment 2 starts away from optimum, making directional selection more relevant there. It is important to note, when the model is run with stabilizing selection only, the phenotypic pliancy and fidelity results are very similar to when evolved with both stabilizing and directional selection (see Supplementary Figure S12, A), but evolution attains a lower fitness in environment 2 (see Supplementary Figure S12, B).

This fitness function selects for individuals for which 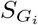 moves closer to 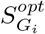. During each generation, reproduction and selection are carried out by setting each simulated cell’s probability of reproduction to be proportional to its fitness. Again, developmentally unstable individuals are considered lethal, thus not included into the next generation. The population of cells reproduces sexually.

Mutation is carried out by allowing each of the nonzero entries of the individual’s gene interaction network, *W*_*GG*_, and gene-environment interaction, *W*_*GE*_ to mutate according to a Gaussian distribution, with mutation rate *μ*. The entries of the PRE matrix, *θ*, are also free to switch between 0 and 1 with a set mutation rate, allowing evolution of susceptibility to PRCs. The mutation rate is set such that during each generation a fraction *μ* of a cell’s *N* genes can be mutated to either become susceptible to PRCs (*θ*_*ij*_ goes from 0 to 1), or lose susceptibility to PRCs (*θ*_*ij*_ goes from 1 to 0). A detailed analysis of the previous model shows robust behavior to a wide range of the model’s parameters [16], which we also observe in parameter testing for our current model (see Supplementary Figure S8 and Supplementary Information).

Note, our computational model is written in Julia (version 1.6.4), and all our model analyses are performed using Julia [57].

### Phenotypic Pliancy Score

Using our computational model, we investigate the degree of phenotypic pliancy upon transferring a population of individuals from the first environmental condition to the second when the PcG mechanisms are intact versus broken. We simulate 1,000 different populations that each evolve under 10 different evolutionary trajectories, where each of these populations is comprised of 1,000 individuals to generate populations with varied intact PcG mechanisms.

At the end of and during the simulated evolution, we measure phenotypic pliancy for each of the 10-million individuals in each population as follows. We consider four environmental conditions: a) the cell is developed in environment 1 and has its polycomb mechanisms intact; b) the cell is developed in environment 2 and has its polycomb mechanisms intact; c) the cell is developed in environment 1, then post-developmentally transferred to environment 2 with intact polycomb-like mechanism(s); and d) the cell is developed in environment 1, then post-developmentally transferred to environment 2 with broken polycomb-like mechanism(s). We only consider those individuals for which the resulting gene expressions are all stable. Each experimental condition then gives rise to a different stable gene expression vector which we denote 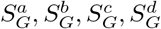, respectively.

We employ the following heuristic to determine when an individual is pliant at gene *i*– if it meets two criteria: 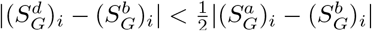 and 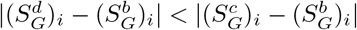. We then calculate the frequency of genes at which that individual is pliant, forming an individual’s pliancy score (see Supplementary Figure S5).

### Descriptions of Head and Neck Single-Cell RNA-Sequencing Data

The entire data set consists of expression data for single cells obtained from 18 patients (5 patients with matching primary and metastatic samples) in two locations: a primary tumor site (oral cavity) with 1,426 cancer cells and 2,817 non-cancer cells, and a metastatic site (lymph node) with 788 cancer cells and 546 non-cancer cells. The non-cancer (normal) cells include fibroblasts, endothelial cells, and B and T cells, amongst others; however, we only consider fibroblasts and endothelial cells. In our analysis, there are four cellular categories: metastatic in lymph node, normal in lymph node, primary tumor in oral cavity, and normal in oral cavity.

### Descriptions of Ovarian Single-Cell RNA-Sequencing Data

The data set consists of expression data for single cells obtained from 9 patients (4 with matching primary and metastatic samples) in three locations and thus cellular categories: a primary tumor site (ovary or fallopian tube) with 1,649 cells, metastatic site (omentum) with 1,062 cells, and normal site (ovary) with 355 cells. The normal cells are comprised of fibroblasts, stromal cells, and mesothelial cells. When we perform PCA on all the cells, the normal cells do not cluster by cell type and the primary cells do not cluster by location.

### Preprocessing of single-cell RNA-sequencing data

Preprocessing and all further analysis of the Puram et. al. and Shin et. al. data sets is performed using the R software package Seurat (version 3.2.3) [48–50]. First, we eliminate cells with fewer than 200 genes expressed in the cellular categories (primary, metastatic, and normal at either both primary and metastatic site or just primary site) and eliminate genes which have non-zero expression in only two cells or less. For H&N data set, 1,426 cancer and 2,817 non-cancer cells from the primary site, and 788 cancer and 546 non-cancer cells from the metastatic site remain. For the ovarian data set, 1,555 cancer and 345 non-cancer cells from the primary site and 1,028 cancer from the metastatic site remain. As for genes after preprocessing, we retain 21,294 genes in the H&N and 18,973 genes in the ovarian data set for further analysis. For H&N data, 65 of these genes are involved in a PcG mechanism including 22 PcG genes, 31 TrxG genes, and 12 genes that have been verified to be controlled by PRCs. For ovarian data, 65 of these genes are involved in a PcG mechanism including 21 PcG genes, 35 TrxG genes, and 9 PRC target genes. We then normalize by log transforming and centering the data and by using SCTransform in the Seurat package [48, 49], finding that the genes’ estimated count-depth relationships were indeed zero for all the cellular categories in each data set. Finally, we regress out cell cycle genes to eliminate any variance due to difference in cell cycles.

### Pliancy z-score for metastatic cells

To measure a metastatic cell’s phenotypic pliancy, we develop a score, referred to as the *pliancy z-score*, that measures the relative extent of movement of a particular metastatic cell’s phenotype toward the normal phenotype and away from the mean of all metastatic cells. This score is calculated as follows. For each gene *g*, let 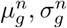 and 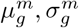 denote the mean and standard deviations of the expression of gene *g* across normal (*n*) and metastatic (*m*) cells, respectively. For each metastatic cell *c*, let *x*_*g,c*_ denote the expression of gene *g* at cell *c* and let 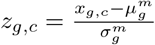 be its *z*-score relative to the distribution across all metastatic cells. Finally let 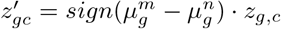. Thus, a positive score means moving away from normal relative to the average, and a negative score means moving toward normal. The cell’s overall score is then devined as 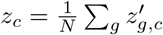, where the sum is taken over the *N* genes included in the analysis.

## Data Availability Statement

The code for both our computational model data generation, model data analysis, and metastatic cancer data analysis can be accessed at: https://github.com/AvivLab/Phenotypic-Pliancy.

## Acknowledgments

The authors would like to thank the reviewers for their time and helpful comments in improving this paper. The authors would like to acknowledge Eduardo Fajardo for help with writing perl scripts for pre-processing the sc-RNA-seq datasets, and Carlos Madrid-Aliste for help with HPC. The authors also would like to thank Moshe Sadofsky, Matthew Scharff, Ruben Coen-Cagli, and Ian Willis for many helpful discussions. Support was provided by NIH CMBG training grant T32-GM007491 to ML, and NIH R01-CA164468 and R01-DA033788 to AB.

## Supporting information

### Methods

#### Model Parameter Testing

We test the sensitivity of the model’s behavior, and the generality of our results, by measuring the change in the percent of cells exhibiting overall phenotypic pliancy when PcG-like mechanism is intact versus broken over a wide range of parameters and 10 randomly chosen starting gene-regulatory network architectures. Specifically, we vary the following five parameters: gene-regulatory network connectivity density (=0.1, 0.3, 0.5), selection strength (=0.5, 1, 2), gene-activation sigmoidal strength (=1, 4, 6), gene threshold level that triggers PRC repression (=0.1, 0.15, 0.2), and mutation rates (gene mutation rate per genome, environment interaction mutation rate, and PcG-like mechanism mutation rate) (=0.1, 0.2, 0.3). Therefore, we obtain results for 243 different parameter settings. We vary these five parameters independently, and we use the same starting population that undergoes the same 100 different evolutionary trajectories for each parameter setting. We then calculate the percent of phenotypically pliant individuals in each evolved population when PcG-like mechanism is left intact or is broken, then calculate the average for each of these two cases over the 100 different evolved populations for each parameter setting. Given that our phenotypic pliancy score (see Materials and Methods) can only be measured for the individuals with broken PcG-like mechanism, we measure overall phenotypic pliancy for each case when PcG-like mechanism is broken vs. left intact to better compare parameter effects. We measure overall phenotypic pliancy by calculating the distance between its stable phenotype after development in its original environment, *S*_*W*_, and the resulting stable phenotype upon transferring it post-developmentally to another environment, 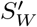 Phenotypic pliancy for a given cell corresponds to a large Euclidean distance 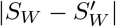, and phenotypic fidelity to a small distance. In our preliminary work, we used a threshold distance of 0.05 to determine phenotypic pliancy, such that if the Euclidean distance is greater than 0.05 then that cell is considered phenotypically pliant. We see a drastic increase in the average phenotypic pliancy when PcG-like mechanism is broken for each of the 243 different parameter settings (p-value = 10^−16^) (see Supplementary Figure 8). Our parameter testing results show the different parameter settings and network architectures do not change our phenotypic pliancy results, strongly suggesting that, while our model lacks biological specificity, it is still biologically relevant in its general implications.

## Supplementary Figures and Tables

**Figure S1.**
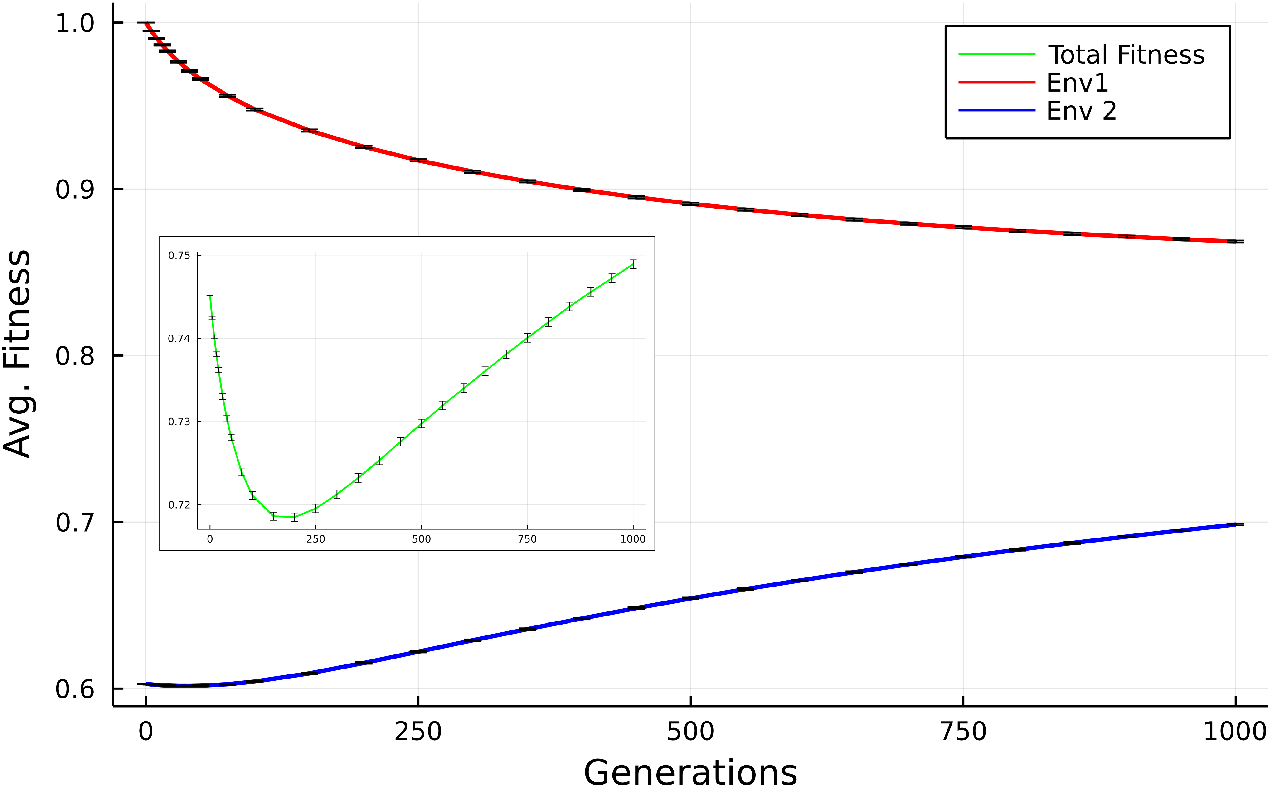
Average Fitness Throughout Evolution: The average combined fitness for both environment 1 and environment 2 shown in the insert due to scale (green), and the average fitness for environment 1 (red) and environment 2 (blue) fitness results.

**Figure S2.**
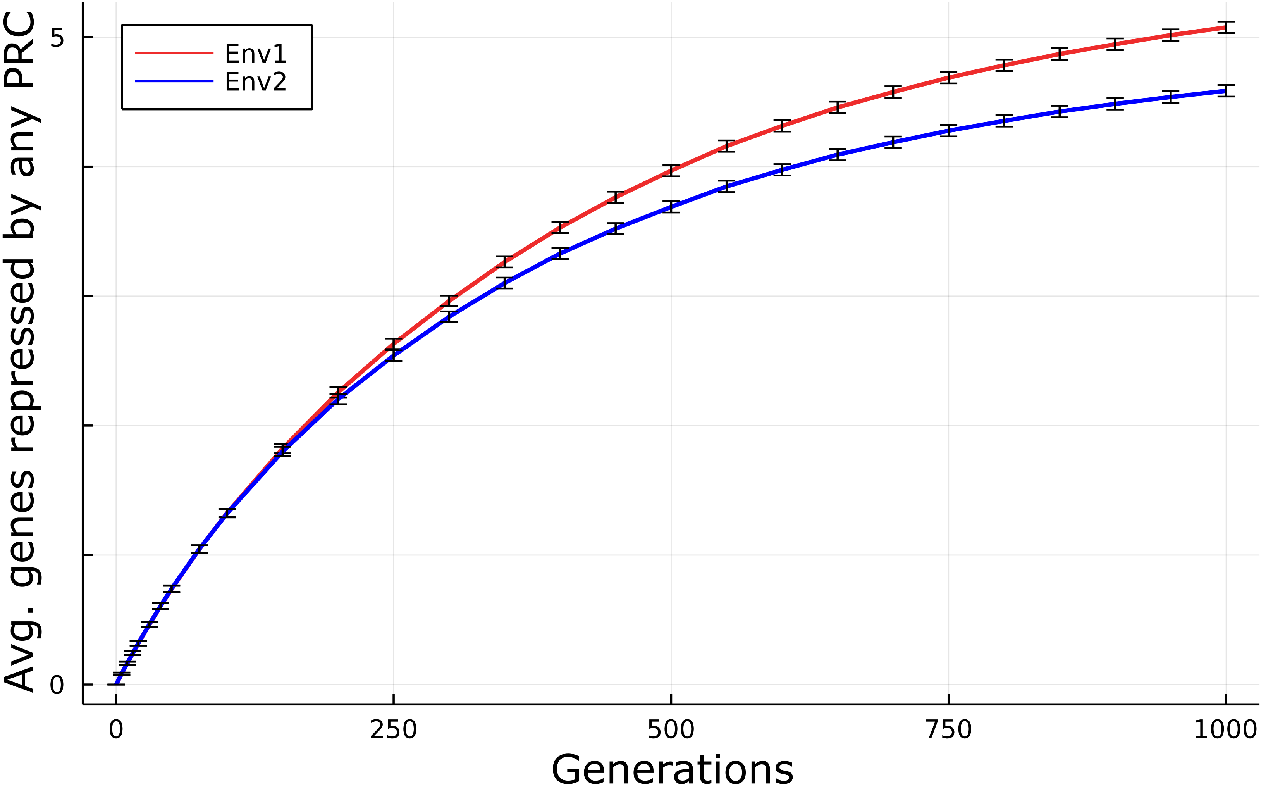
Genes Repressed by PcG-like mechanisms During Evolution: The average number of genes that are repressed by any PRC during evolution in environment 1 (red) or environment 2 (blue).

**Figure S3.**
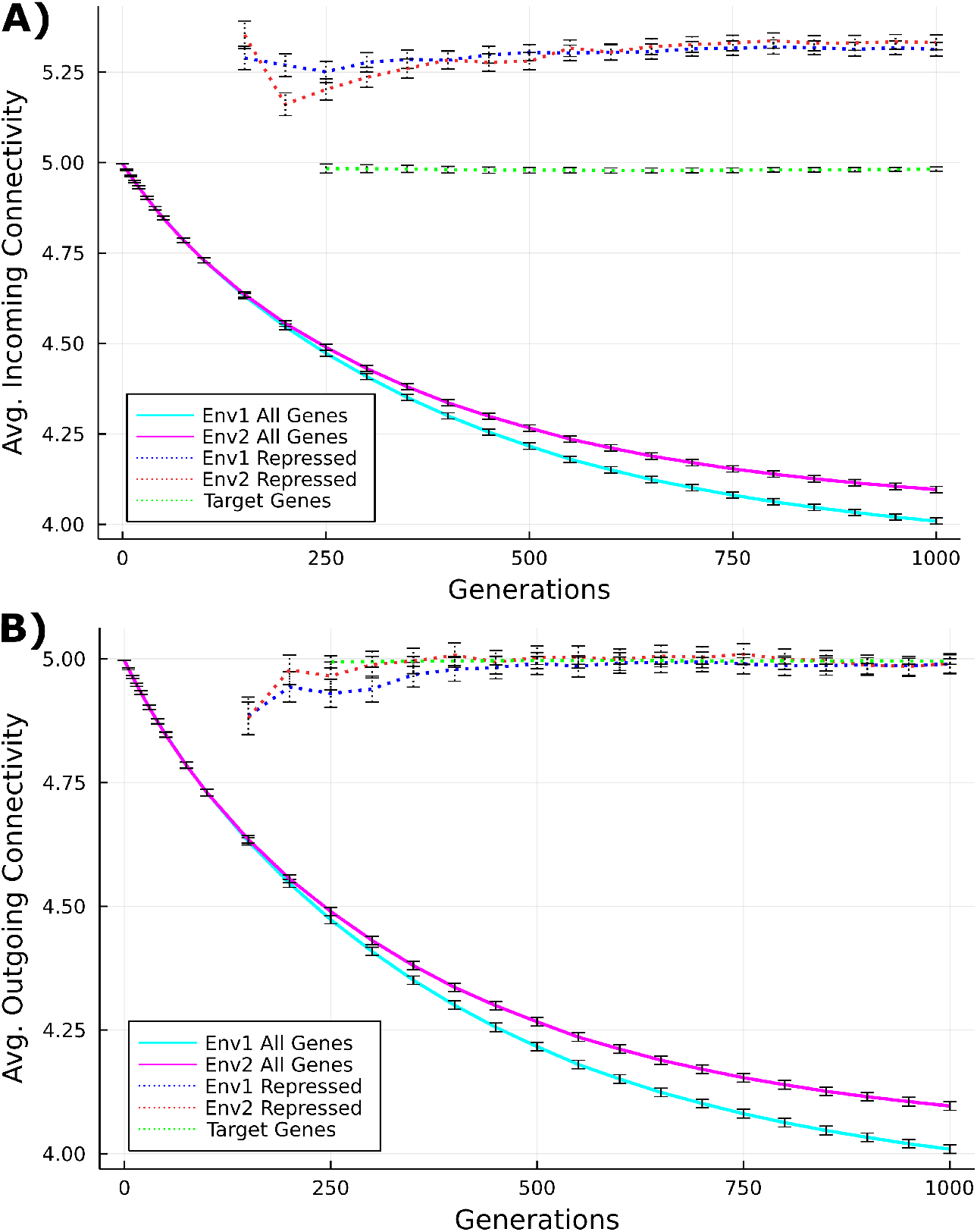
Connectivity of the Gene Regulatory Networks Throughout Evolution: The average incoming (A) and outgoing (B) effective connectivity of the gene regulatory networks for all 10 million cells (solid lines). The effective connectivity averaged across all the genes in the network for environment 1 and environment 2 decreases throughout evolution as would expect (magenta and cyan solid lines). The average connectivity of the target PRC genes, whether repressed or not during development (green dotted lines), and only the genes repressed by any PRC (red and blue dotted lines) are shown as well. Genes that are repressed by any PRC have a higher average incoming connectivity as compared to all the target genes. Note, we look at average connectivity, as opposed to average effective connectivity, for PcG-like mechanism target genes that are repressed or not to show the connections the repressed target had before repression.

**Figure S4.**
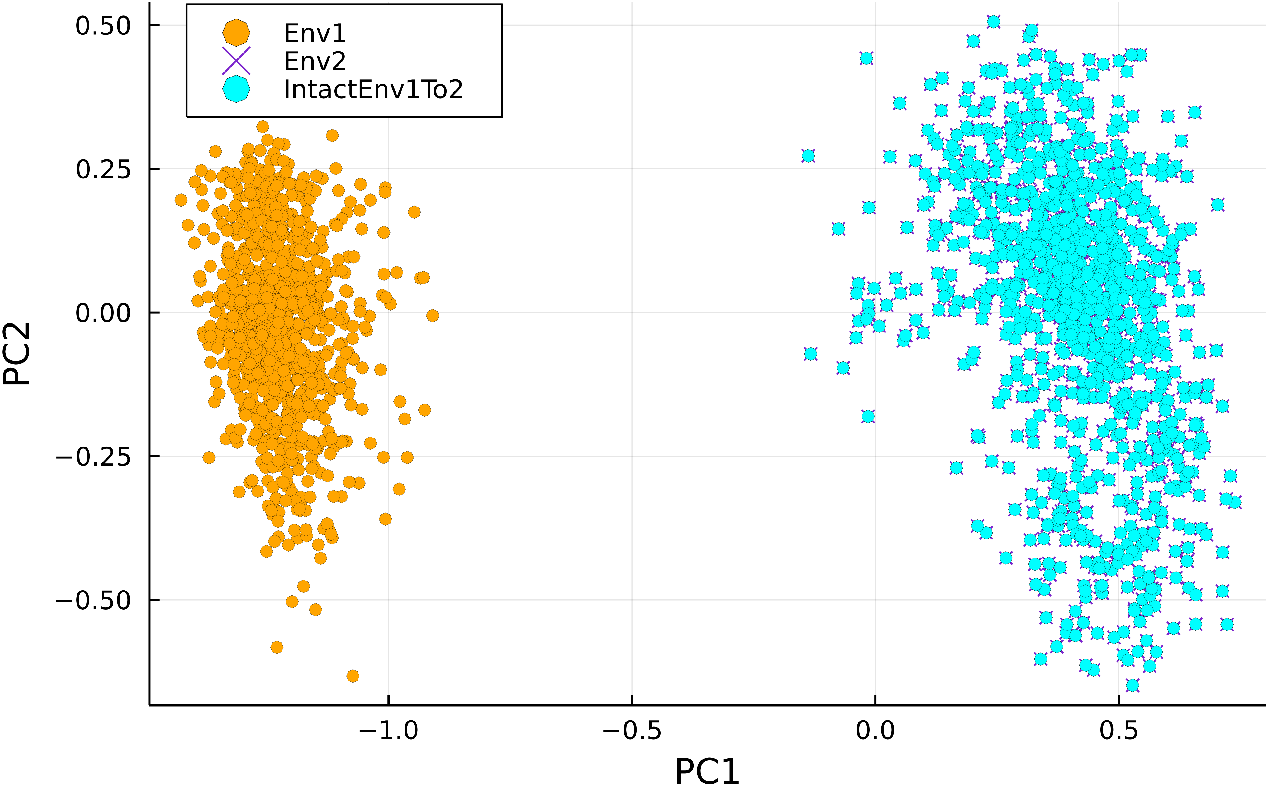
Phenotypic Pliancy and Fidelity Results from Computational Model without the evolution of PcG Mechanisms: PCA results for all individuals in population at the end of evolution (after generation 1,000) when evolve with no PcG mechanism evolution to assess phenotypic fidelity. After evolution when transfer from environment 1 to environment 2 (cyan) and compare to environment 1 (orange) and environment 2 (purple X’s), we see that the individuals move from environment 1 to environment 2 and thus do not exhibit phenotypic fidelity like they do when evolve with polycomb mechanism. Note, when evolve without polycomb and move from environment 1 to environment 2 (cyan), the results exactly overlap with environment 2 (purple X’s) because the networks for these cases are the same when do not have any polycomb repressed genes.

**Figure S5.**
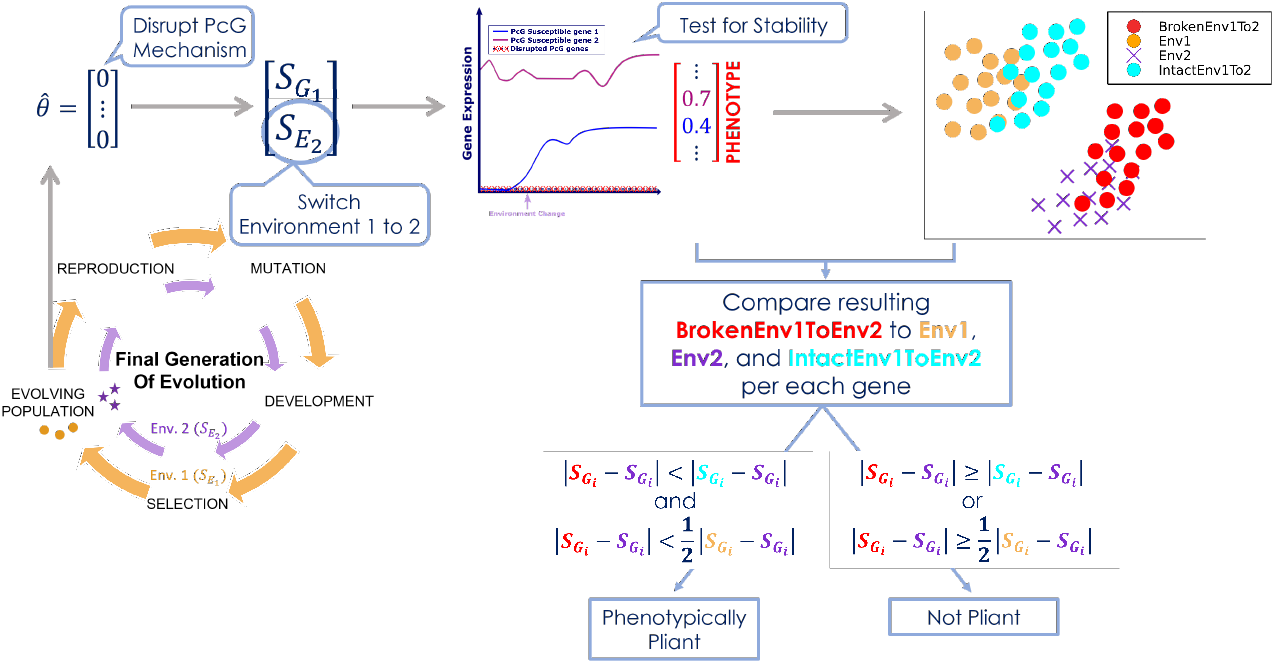
Schematic for Measuring Phenotypic Pliancy in the Model: We measure phenotypic pliancy of a simulated cell by first disrupting the PcG mechanism by removing its control from its repressed genes, we then switch the cell from environment 1 (orange) to environment 2 (purple), we test for stability of the gene expression pattern in this new environment, and finally we determine if this resulting stable gene expression pattern, or phenotype, when break PcG and switch to environment 2 (red) is phenotypically pliant. Mathematically, we first compare if the red moves closer to environment 2 in purple than environment 1 in orange, and then compare if the red is closer to environment 2 in purple than when PcG is intact but switched environments in cyan is to environment 2 in purple. If these conditions are true, then that gene is considered considered phenotypically pliant. Finally, a cell’s phenotypically pliancy score is the percent of genes that move closer to the environment switched to when PcG is broken (red) but remain closer to original environment when PcG is left intact (cyan).

**Figure S6.**
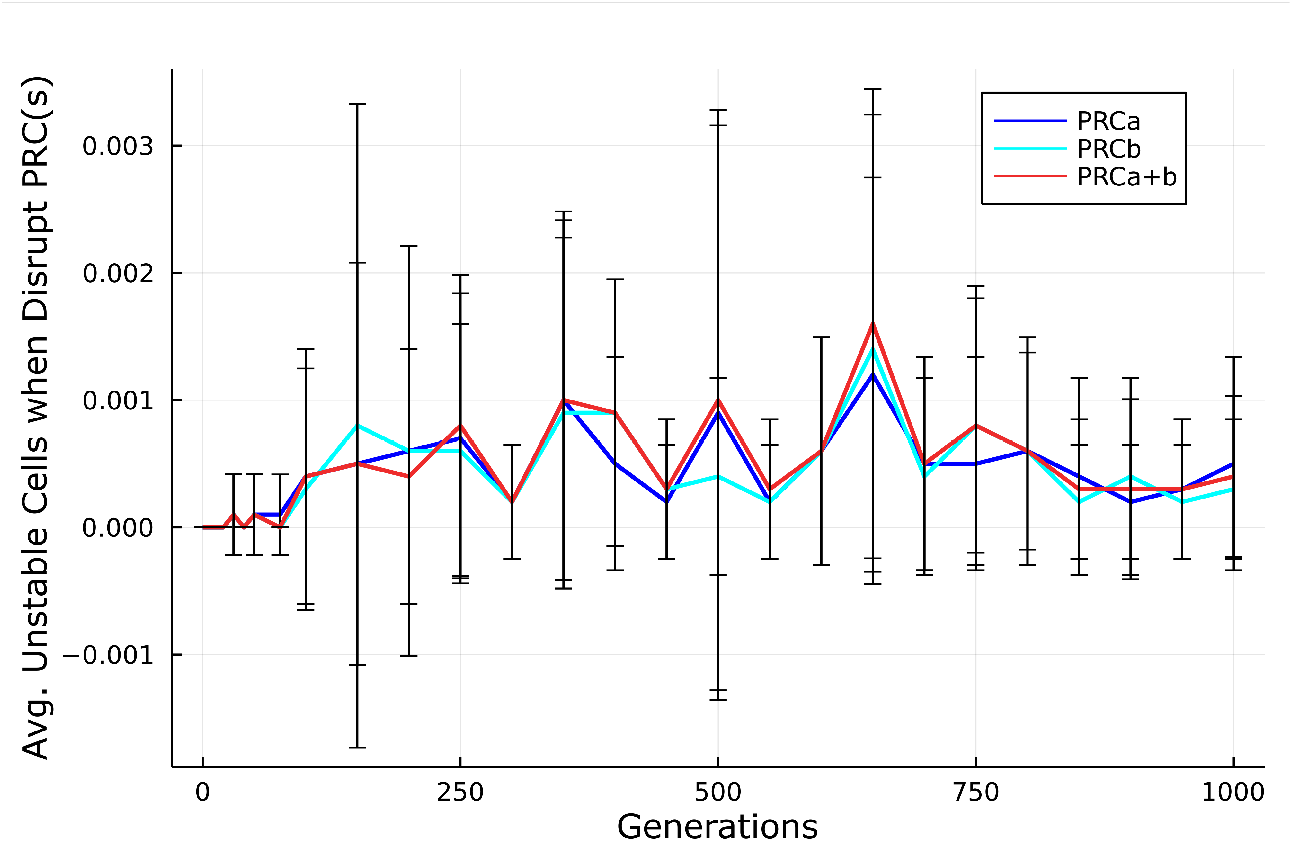
Average Number of Unstable Cells For Varying Degree of PcG-like Mechanisms Breakage: The average number of unstable cells in each of the 10,000 different populations when break PRCa alone (blue), PRCb alone (cyan), and PRCa and PRCb together (red).

**Figure S7.**
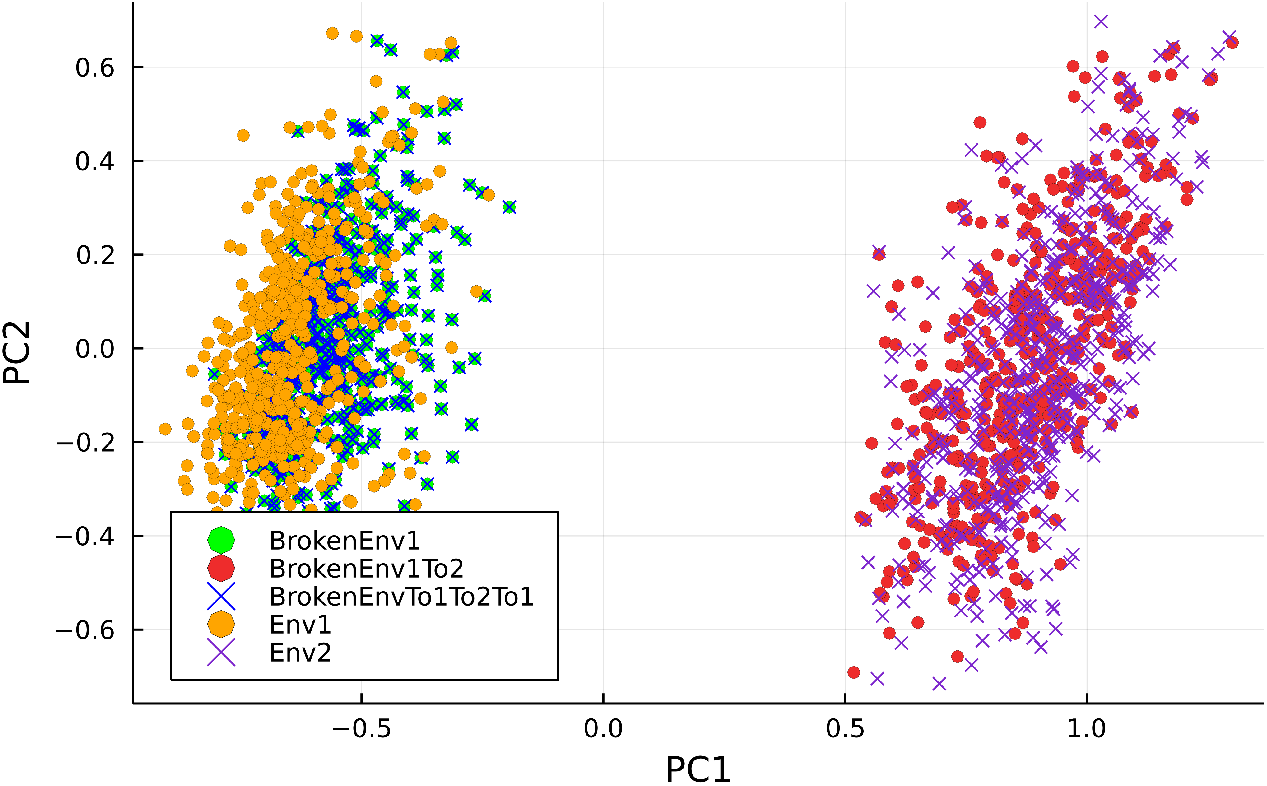
Sustained Phenotypic Pliancy due to PcG-like Breakdown in Model: Principle Component Analysis (PCA) showing that when PcG-like mechanisms are dysregulated in our model that phenotypic switching is sustainable, such that after switching from environment 1 to environment 2 (red circles) and then back to environment 1 (blue X’s) that the phenotypes more closely resemble what environment they were switched to. As a control, we also see that if break PcG-like mechanisms but keep in environment 1 (green circles) then phenotype stays close to that of the evolved phenotype in environment 1 (orange circles).

**Figure S8.**
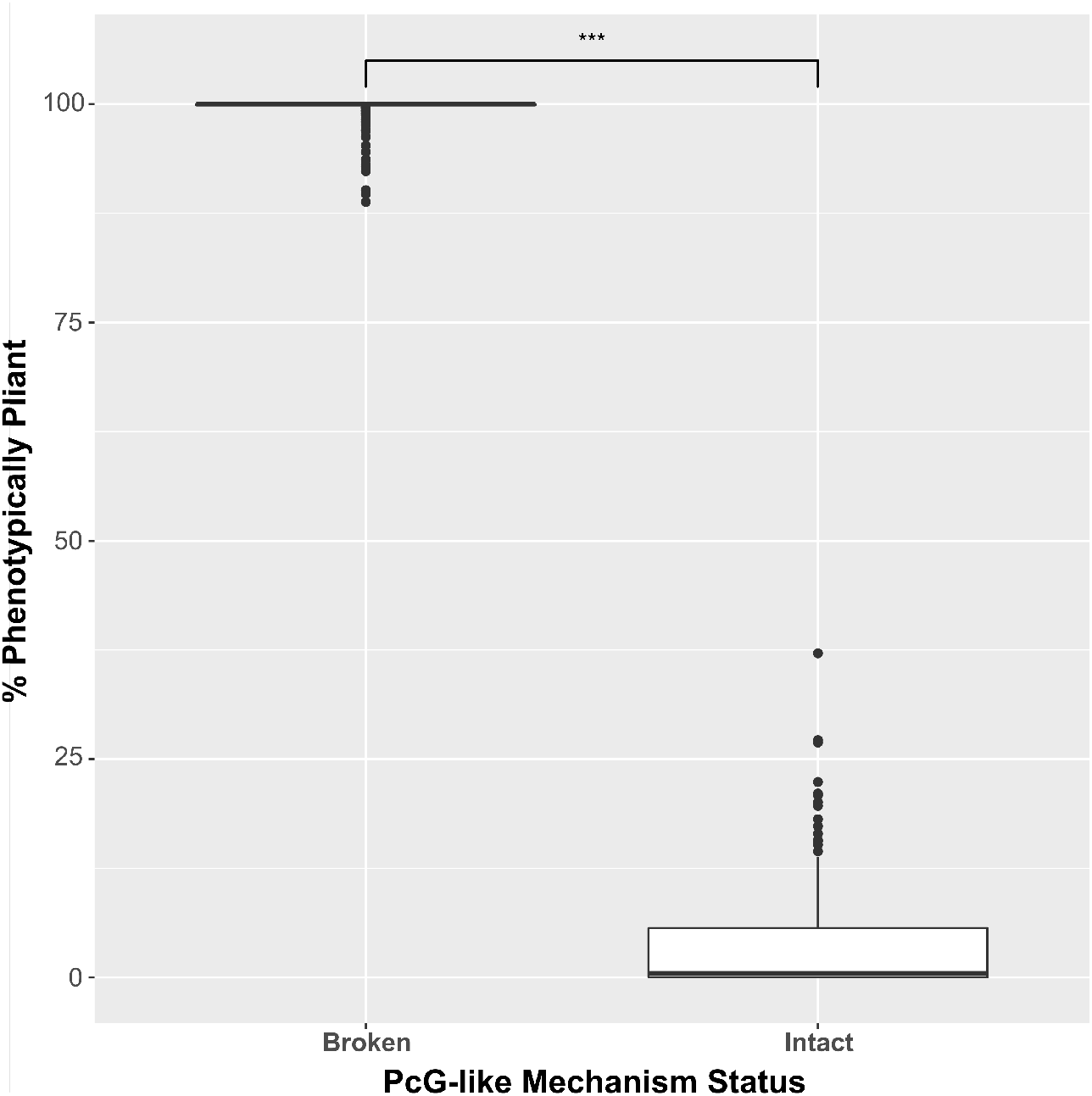
Overall Phenotypic Pliancy for Model Parameter Testing: Phenotypic pliancy when PcG-like mechanism is intact versus broken over a wide range of parameters and 10 randomly chosen starting gene-regulatory network architectures. The percent of cells that are phenotypically pliant when PcG-like mechanism is broken is statistically significantly greater than when the PcG-like mechanism remains intact (p-value *<* 0.001).

**Figure S9.**
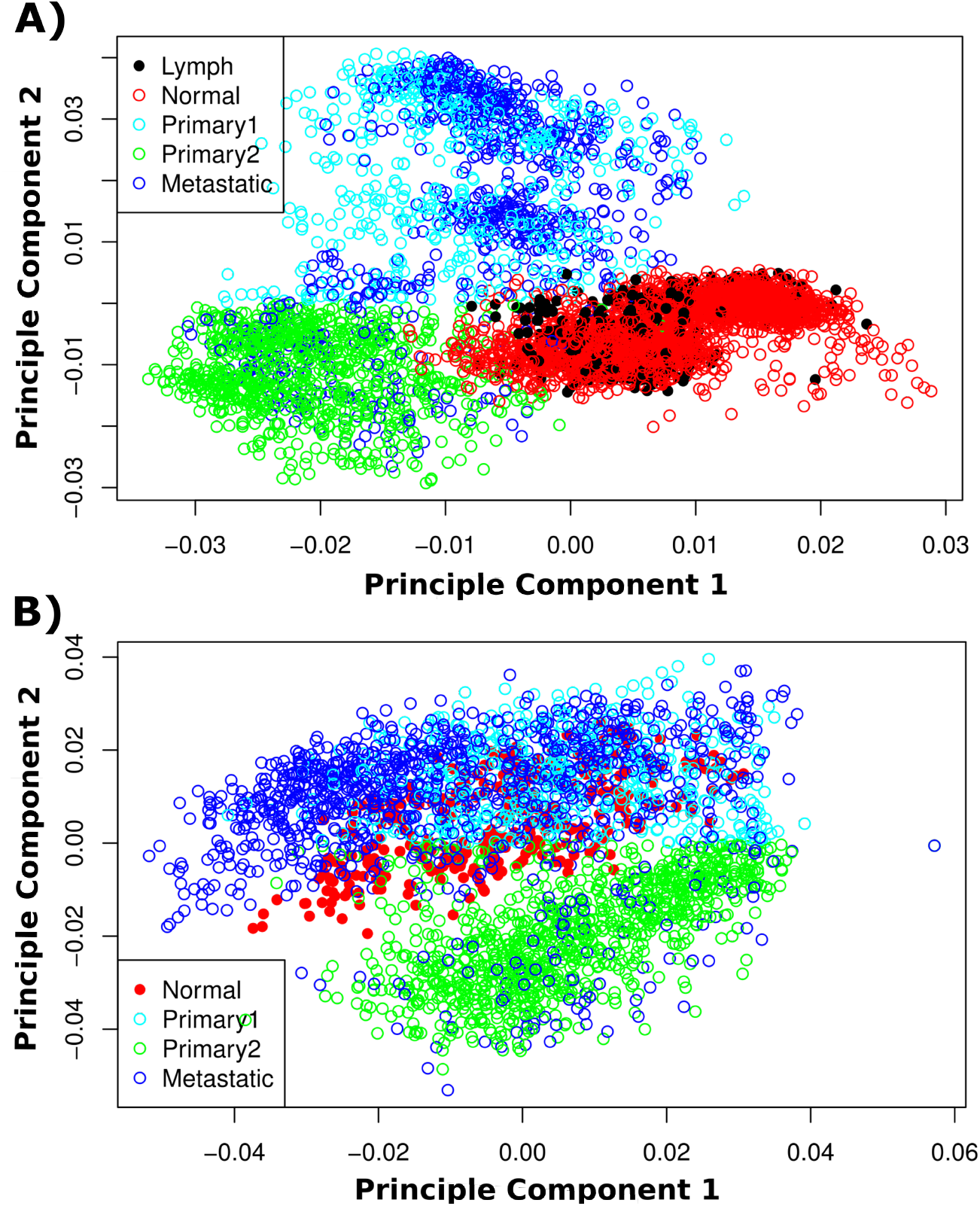
PCA Results for Metastatic Cancer Datasets: Principle Component Analysis (PCA) results for **A**. Head and Neck and **B**. Ovarian metastatic cancer dataset, where metastatic cells’ phenotypes (given by their gene expression patterns) are each represented by dark blue circles, primary cancer cell’s phenotypes are represented by both green and cyan, and the normal non-cancer cells’ phenotypes at the metastatic site and primary site are represented by black and red circles, respectively. Note, the primary cells are split by principle component 2 (y-axis), such that primary cells with PC2 values greater than zero are represented by cyan circles and primary cells with values less than zero are represented by green circles.

**Figure S10.**
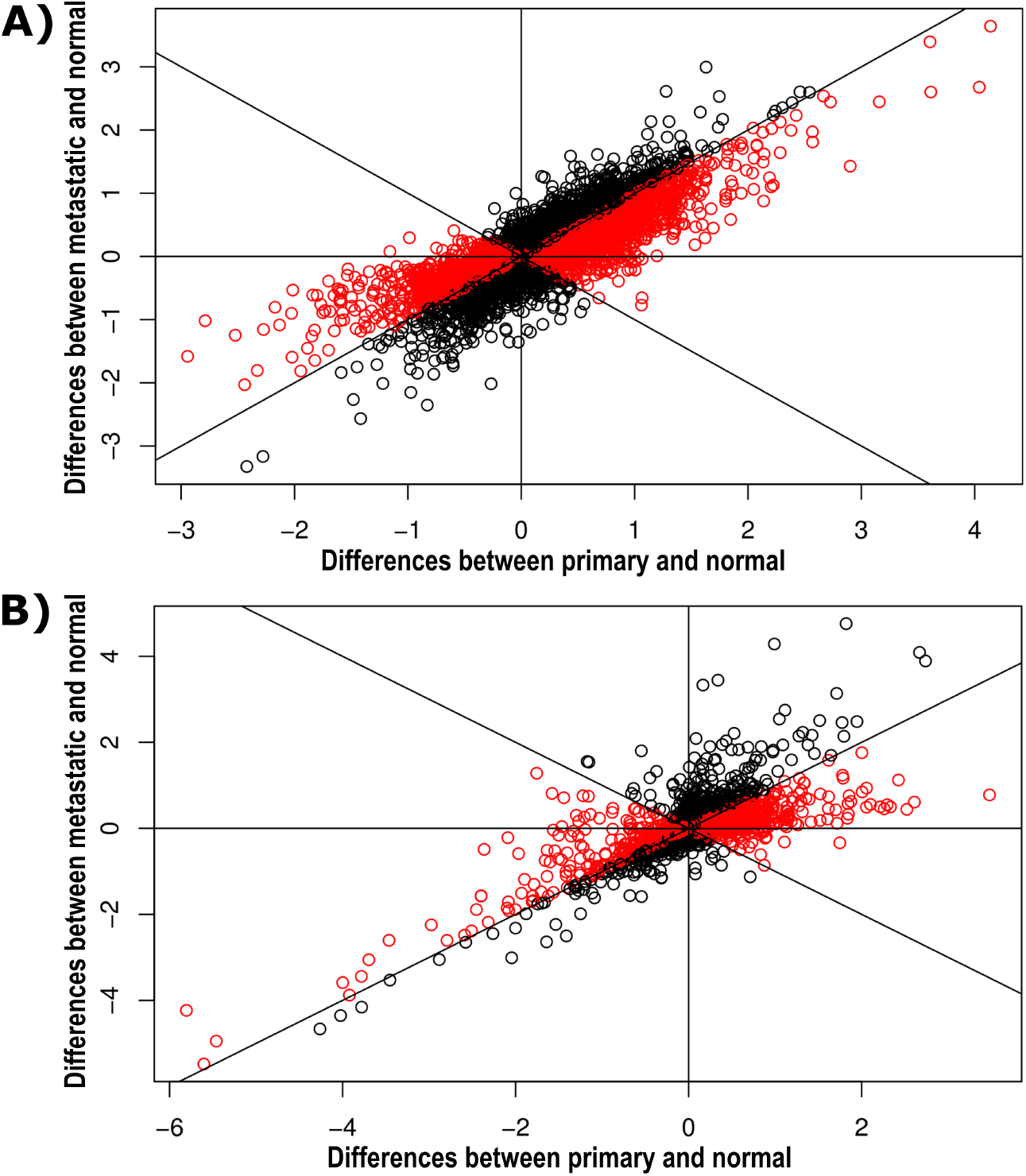
Gene-Wise Differences for Metastatic Cancer Datasets: Plot of the gene-by-gene differences between primary and normal at the primary site (x-axis) versus the differences between metastatic and normal at the lymph node site or ovary site for the **A**. head and neck and **B**. ovarian metastatic cancer dataset, respectively (y-axis). We have included lines *y* = *x* and *y* = −*x* for comparison. There is a trend for the gene-wise differences between metastatic and normal at the lymph node site being smaller in absolute value than gene-wise differences between primary and normal at the primary site, which is the case for **A**. 68% of the genes differentially expressed between primary tumor versus metastatic cells (p-value *<* 10^−8^) for H&N (shown in red), and **B**. 72% (p-value *<* 10^−9^) for ovarian (shown in red).

**Figure S11.**
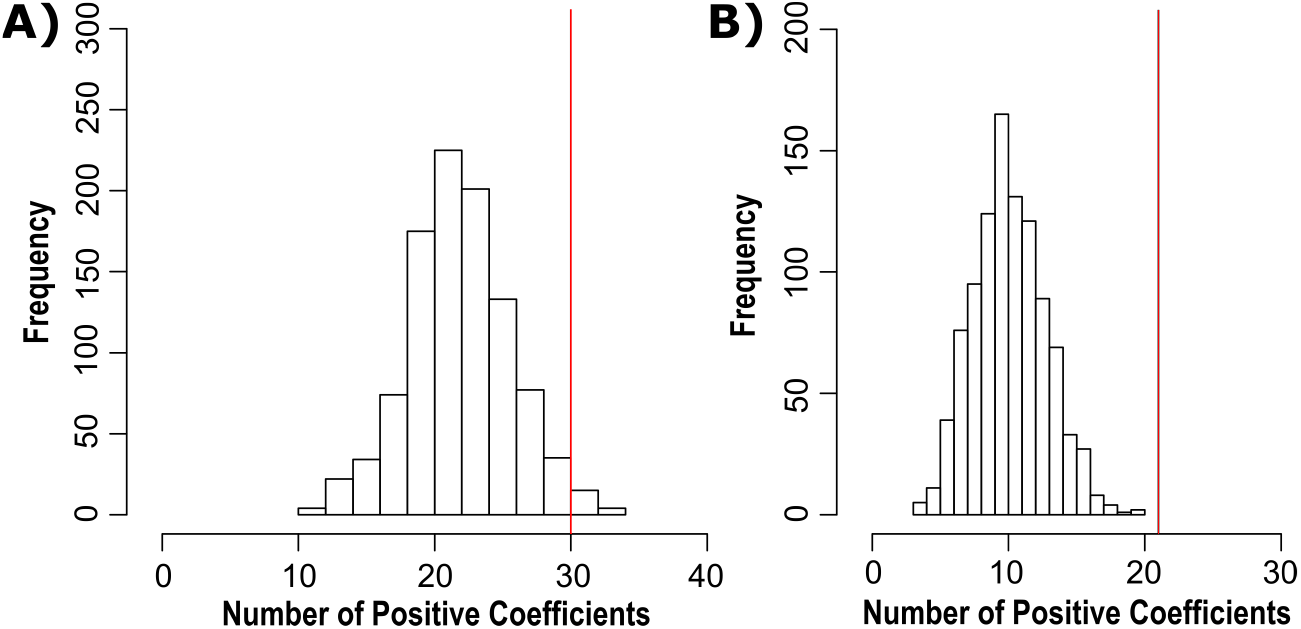
Significance Assessment of PcG Genes as Source of Pliancy: Histogram of the bootstrapping results, which we perform by randomly picking 1000 samples of PcG mechanism genes and performing linear regression of the score as a function of these genes’ expression profiles for **A**. H&N and **B**. ovarian data, respectively. Each plot is the distribution of the positive coefficients computed for each random sample. The red line is the number of PcG mechanism genes with positive coefficients, which is 30 for H&N data and 21 for ovarian data. The mean is 22.3 out of 53 for H&N data, and mean is 10.6 out of 56 for the ovarian data, so the mean for both **A**. and **B**. (red line) is statistically significant (p-value *<* 0.05 and p-value *<* 10^−5^, respectively).

**Figure S12.**
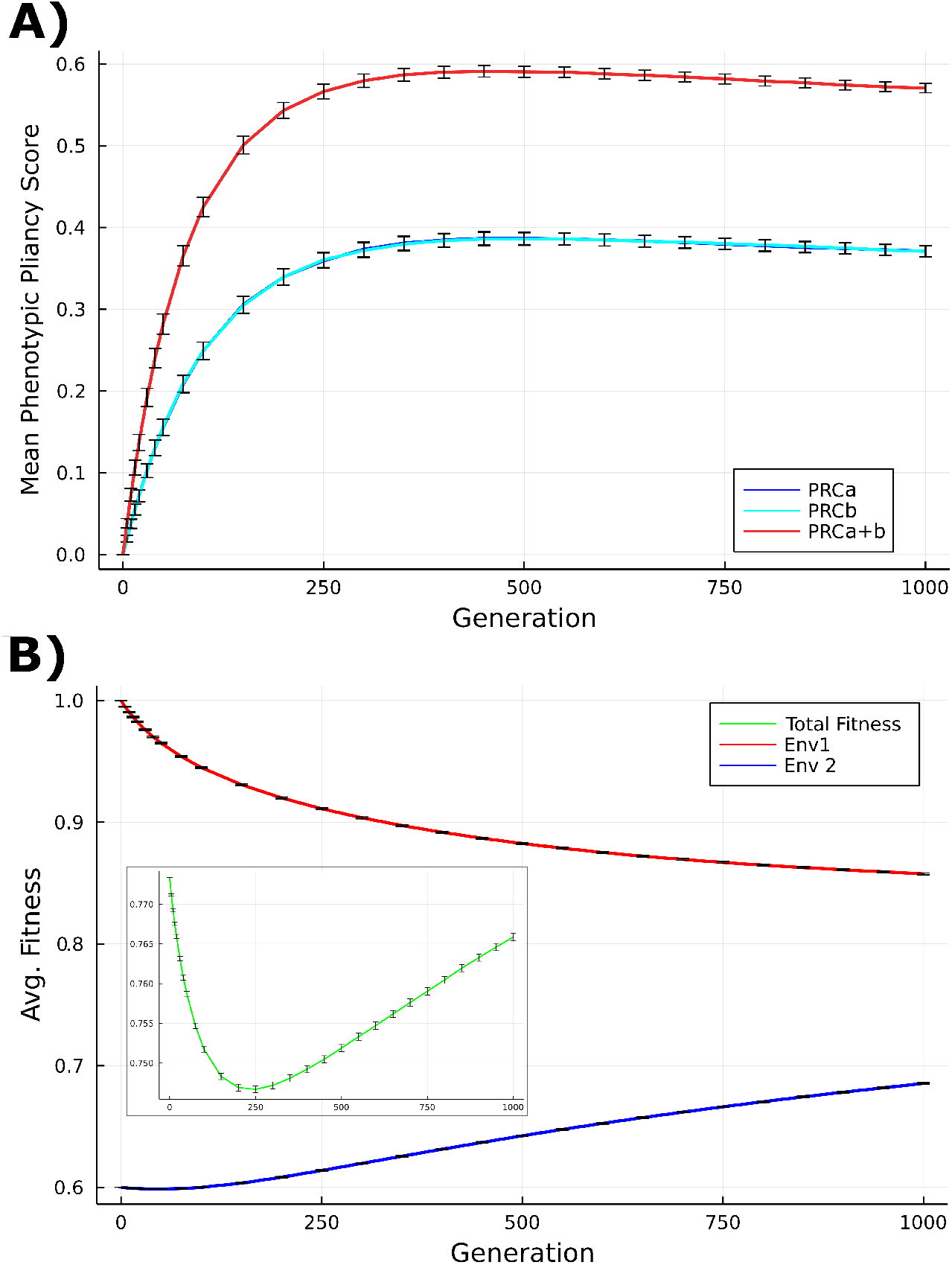
Phenotypic Pliancy and Fitness Results when Evolve with Stabilizing Selection Only: **A**. Average phenotypic pliancy score when vary degree of PcG mechanism dysregulation during evolution for all 10,000 populations when move from environment 1 to environment 2 to quantitatively assess pliancy when evolve with stabilizing selection only. We vary degree of dysregulation by breaking PRCa alone (blue), PRCb alone (cyan), or both PRCa and b (red) for all 10,000 populations for a total of 10 million simulated cells pliancy score averaged (y-axis) for different generations throughout evolution (x-axis). **B**. The average combined fitness for both environment 1 and environment 2 shown in the insert due to scale (green), and the average fitness for environment 1 (red) and environment 2 (blue) fitness results when evolve with stabilizing selection only.

**Table 1.** Polycomb Mechanism Genes Used in Single-Cell RNA-Sequencing Data Analysis:

| PcG Protein | TrxG Protein | Controlled by PRC |
| --- | --- | --- |
| RING1 | CHD8 | PCDH15 |
| RNF2 | ASH2L | PCDHB1 |
| CBX2 | SMARCA1 | PCDHB4 |
| CBX4 | SMARCA2 | PCDHB15 |
| CBX6 | RBBP5 | CDH8 |
| CBX7 | WDR5 | CDH13 |
| CBX8 | CHD3 | CDH18 |
| PCGF1 | CHD4 | CDH19 |
| PCGF2 | CHD1 | CDH23 |
| PCGF3 | CHD2 | RAP1A |
| BMI1 | MTA1 | RAP1B |
| PCGF5 | MTA2 | RAP1GAP |
| PCGF6 | MTA3 |  |
| SCMH1 | HDAC1 |  |
| RYBP | HDAC2 |  |
| YAF2 | MBD2 |  |
| EZH1 | MBD3 |  |
| EZH2 | POLD3 |  |
| EED | BPTF |  |
| SUZ12 | SMARCB1 |  |
| RBBP4 | DPF1 |  |
| RBBP7 | KMT2A |  |
|  | KMT2D |  |
|  | KMT2C |  |
|  | KMT2B |  |
|  | KAT8 |  |
|  | DPY30 |  |
|  | SETD1A |  |
|  | SETD1B |  |
|  | CXXC1 |  |
|  | WDR82 |  |
|  | KDM6A |  |
|  | NCOA6 |  |
|  | PAXIP1 |  |
|  | PAGR1 |  |

**Table 2.** Differential gene expression analysis for metastatic cancer data sets.

| Data (Tumor) | DE PcG Genes | Log-Fold Change Sum |
| --- | --- | --- |
| H&N (Primary) | 42 genes | -20.1 |
| H&N (Metastatic) | 45 genes | -29.2 |
| Ovarian (Primary) | 25 genes | 20.2 |
| Ovarian (Metastatic) | 16 genes | -14.3 |

**Table 3.** Model Parameters and Values.

| **Parameter** | **Value** |
| --- | --- |
| Genes | 50 |
| Population Size | 1,000 |
| Gene Regulatory Network Connectivity | 0.1 |
| Generations | 1,000 |
| Environment Components | 50 |
| Maximum Iterations | 100 |
| Gene Mutation Rate | 0.1 |
| Environment Interaction Mutation Rate | 0.1 |
| PcG-like Mechanism Mutation Rate | 0.1 |
| Selection Strength | 0.5 |
| Gene-Activation Sigmoidal Strength | 1.0 |
| PRC Repression Gene Threshold Level | 0.15 |
| PRC Critical Time Point | 2 |
| Number Different Environments | 2 |
| Number of Different PRCs | 2 |
| Proportion of Env. Components Affecting Cell > 0 | 0.4 |
| Proportion of Genes Able to be Affected by Envs. | 0.4 |
| Proportion of Env. Components Affecting Gene | 0.4 |
| Minimum Difference Between Env. 1 and Env. 2 | 70% |
| Minimum Difference in Env. Optimum States | 40% |
| Individual Weights | Gaussian |
| Sexual Reproduction Flag | true |

## Notes

https://github.com/AvivLab/Phenotypic-Pliancy

### Competing Interest Statement

The authors have declared no competing interest.

### Summary of Updates

Added control experiments as well as reorganizing the manuscript order

